# Multi-body Fluctuation-Induced Forces Between Membrane Proteins: Insights from Mesoscale Simulations

**DOI:** 10.1101/2025.09.12.675822

**Authors:** Adrià Bravo Vidal, Weria Pezeshkian

## Abstract

The spatial organization of membrane-associated proteins is essential for a wide range of cellular processes, including signal transduction, endocytosis, and cell adhesion. While protein clustering can be driven by direct short-range forces, indirect interactions mediated by the membrane itself, particularly those arising from thermal shape fluctuations, are potentially sufficient to drive clustering in the absence of direct binding. In this study, we investigate how fluctuation-induced interactions contribute to the lateral organization of membrane inclusions using mesoscale simulations. Our approach is based on dynamically triangulated surfaces and is parameterized by three mesoscale quantities that capture local membrane rigidification and curvature induction. We show that local membrane rigidification drives the non-random organization of membrane inclusions and, above a critical concentration threshold, induces a fully segregated state. This threshold depends strongly on the magnitude of the induced rigidification. We further demonstrate that membrane tension only weakly affects lateral organization away from the threshold but has a pronounced effect near it. Extending our analysis to spherical geometries, we obtain similar behavior relevant to experiments on small unilamellar vesicles. In mixed systems containing two types of stiff inclusions, we find that stiffer proteins act as nucleation centers for softer proteins. Finally, we show that protein-induced curvature, combined with fluctuation-mediated clustering, can drive membrane shape remodeling. Our results are consistent with previous findings while additionally extending the characterization across the full parameter space and to new conditions of direct biological relevance. Overall, our findings suggest that local suppression of membrane-shape fluctuations by proteins generates effective attractive forces capable of driving protein reorganization on membranes, with broad implications for cell biology and the design of membrane-associated nanoparticles.

**STATEMENT OF SIGNIFICANCE:** This study demonstrates that membrane protein lateral organization can emerge solely from membrane-mediated interactions driven by thermal shape fluctuations, independent of direct protein–protein interactions. Although theoretical models have long predicted such forces, their non-additive nature and apparent weakness left it unclear whether they could overcome mixing entropy. Here, we show that these forces can indeed be sufficient to drive clustering and, crucially, that they promote the formation of heterogeneous protein assemblies. These findings suggest that protein accumulations observed in experimental membrane-imaging data may arise from membrane fluctuations rather than direct affinities, highlighting the need for careful consideration when interpreting membrane-imaging experiments.

## I. INTRODUCTION

Many proteins must cluster and localize on the membrane surface to carry out specific functions (1, 2). This spatial organization is critical for various cellular processes such as signal transduction, endocytosis, and cell adhesion (3). Understanding how and why proteins organize on membranes is therefore essential to uncovering the mechanisms behind many vital cellular processes (4). Membrane-associated proteins can cluster through both direct and indirect interactions (5). Direct forces, such as electrostatic and van der Waals interactions, are typically short-range. Indirect interactions, however, emerge from the membrane’s physical response to perturbations caused by embedded or adhered proteins. These membrane-mediated forces include short-range effects like hydrophobic mismatch and capillary interactions (6, 7), as well as longer-range forces arising from perturbations on membrane shapes, such as local curvature or suppression of thermally induced shape fluctuations. The latter, which are the main focus of this article, are particularly compelling due to their long-range and generic nature, rendering them versatile mediators of cellular function. These interactions are also exploitable by viruses and toxins, and they also hold promise for the rational design of drug delivery systems. For example, it is hypothesized that Shiga toxin clusters on the membrane via fluctuation-induced forces to facilitate its cellular uptake (5, 8).

Over the past few decades, numerous theoretical studies have investigated the behavior of membrane-mediated interactions arising from perturbations in the membrane shape (9–13). However, it is not fully clear whether these interactions alone can lead to protein clustering, especially given that these forces are non-additive and on the order of the thermal energy *k*_*b*_*T* (8, 14). These forces can be classified into two types based on the perturbations induced by proteins on membranes and their physical characteristics; (i) curvature-mediated forces, arising from the ability of proteins to induce local membrane curvature and (ii) fluctuation-induced forces, originating from the capacity of proteins to suppress or alter membrane fluctuations, and emerging even in cases where no local curvature is induced.

For two disk-like, stiff proteins that bind tightly to the membrane, at large distances (much greater than the protein size), the leading term of the fluctuation-induced interaction as a function of the inter-protein distance (*r*)decays as 1*/r*^4^ in bending-dominated regimes, and as 1*/r*^8^ in tension-dominated regimes (9, 12). Higher-order corrections do also exist (13) and the interaction is non-additive: the total force in a many-particle system cannot be obtained by simply summing pairwise interactions. Neither non-additivity nor higher-order corrections are unique to fluctuation-mediated forces; similar effects are well known in physics more broadly, including depletion and van der Waals interactions (15–18). However, for fluctuation-mediated forces in biologically relevant regimes, i.e., at separations comparable to only a few protein diameters, higher-order corrections and genuine many-body interactions become comparable or even greater in magnitude to the leading asymptotic term. For example, Yolcu et al. (13), show that the two-body interaction can differ by nearly an order of magnitude from its leading asymptotic approximation at these distances.

Consequently, effective models of membrane protein organization must explicitly account for membrane shape fluctuations and cannot, in general, be reduced to additive pairwise interactions, especially using only the asymptotic contribution. Therefore, determining whether collective membrane-mediated attractions are sufficient to overcome mixing entropy and drive clustering cannot be resolved from pairwise interactions alone. However, the resulting many-body problem is analytically intractable in most cases, making computational approaches essential.

In recent years, much progress in mesoscale modeling of biological membranes has been made (19, 20) that allows for exploring multi-body membrane-mediated interactions (21–23). For rotationally symmetric proteins that bind and unbind to the membrane locally increasing its stiffness, a first-order phase transition between a dispersed and an aggregated state has been predicted to take place when the proteins are sufficiently rigid (24–26). Similarly, aggregation of membrane-bound, curvature-inducing stiff proteins was observed as a consequence of protein-induced stiffness (27, 28), which could even lead to the formation of membrane invaginations (29). Despite these advances, our understanding of fluctuation-induced forces remains incomplete, and many questions are still unanswered, making it difficult to evaluate their relevance in biological contexts. For instance, it remains unclear how fluctuation-induced forces behave in complex systems, such as membranes containing more than a single protein, or how key physical parameters, like membrane tension and spontaneous curvature, influence the clustering. This extends beyond the intrinsic properties of the membrane to the features and strength of protein–membrane interactions. In particular, the role of protein-induced changes in the Gaussian modulus in a many-body context has not yet been explored.

Last but not least, while curvature-mediated interactions on intrinsically curved membranes (e.g., small vesicles) have been studied (30–32), the corresponding role of fluctuation-mediated forces in such geometries remains largely unaddressed.

In this work, we employed mesoscale simulations based on the dynamically triangulated surface (DTS) model to systematically investigate the role of fluctuation-induced forces in the clustering of rotationally symmetric membrane proteins, which represent a large class of proteins at the mesoscale (33–35). Specifically, we examined how protein rigidity (i.e, the rigidity induced on the membrane upon binding), protein concentration, and membrane tension influence clustering in both planar and vesicular geometries. Going beyond simple systems, we also studied membranes containing two distinct protein types and the impact of protein-induced spontaneous curvature. While our results recover known behaviors, they also uncover previously unreported features, highlighting the rich and nuanced nature of fluctuation-mediated interactions. Ultimately, we provide a clear parameter-based framework for understanding protein clustering driven by large-distance, membrane-mediated interactions.

## II. METHODS

### A. Dynamically triangulated surface simulations

At length scales much larger than its thickness, a lipid membrane can be effectively modeled as a two-dimensional surface embedded in three-dimensional space. Several computational methods take advantage of this property to describe membrane shape and behavior (20, 36, 37). In this work, we used the FreeDTS software, which enables the simulation of a wide range of membrane-involved biological processes at the mesoscale (23). This model has been well validated and reproduces theoretically tractable thermodynamic properties of membranes, such as the characteristic undulation spectrum for flat membranes and mechanical energy of vesicles (23, 38–44). We furthermore verify that the model correctly reproduces the undulation spectrum of vesicles (see Fig. S1). In FreeDTS, a membrane is represented by a dynamically triangulated, self-avoiding mesh comprising *N*_*ν*_ vertices, *N*_*l*_ edges and *N*_*t*_ triangles that satisfy the Euler characteristic *ξ* = *N*_*ν*_ − *N*_*l*_ + *N*_*t*_. Self-avoidance is enforced by rigid spherical beads of diameter *l*_DTS_ at each vertex, a maximum edge length between neighboring vertices of 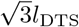 and a mild dihedral angle constraint between neighboring triangles. The constant *l*_DTS_ is set as the unit length and can be converted to physical units by comparison to the biological system under study, as specified below. The energy associated to the membrane shape is represented by the Helfrich Hamiltonian (45), which goes up to second order in surface curvatures. Its discrete formulation takes the form

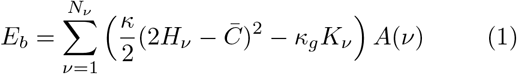

where 2*H*_*ν*_ = *c*_1_(*ν*) + *c*_2_(*ν*) and *K*_*ν*_ = *c*_1_(*ν*)*c*_2_(*ν*) are the mean and Gaussian curvature associated with vertex *ν*, and *κ, κ*_*g*_ and 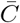 are the bending rigidity, Gaussian modulus and spontaneous curvature of the bilayer. The sum goes over all vertices. Note that here the Gaussian curvature term is written with a minus sign, so *κ*_*g*_ and its local changes are reported as positive quantities, whereas in much of the literature a positive sign is used, in which case *κ*_*g*_ is reported as negative. Here, we focus on symmetric bilayers; thus, 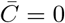. For more details, see Supplemental Note 1. Various methods exist to calculate geometric properties associated with each vertex *ν* on a triangulated surface (46–48). We employ a method based on the shape operator approach implemented in FreeDTS (38), which uses a discretized shape operator to obtain the vertex normal, **N**_*ν*_, surface area *A*(*ν*), and principal curvatures *c*_1_(*ν*) and *c*_2_(*ν*), along with their associated principal directions **T**_1_(*ν*) and **T**_2_(*ν*).

In addition to the bending energy, different energy contributions might be added to include different constraints. To keep the total surface area of the membrane constant, the energy is coupled to a second-order surface-area potential 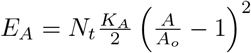, where *A*_*o*_ is the desired surface area and *K*_*A*_ is the compressibility modulus. For membrane patches subject to PBC in the XY direction and constant frame tension *τ*, the projected area *A*_*p*_ of the patch is coupled to an energy of the form *E*_*τ*_ = − *τ A*_*p*_. For closed geometries, changes in volume are energetically penalized through a coupling to the enclosed volume, *E*_*V*_ = − Δ*PV*, where Δ*P* represents the pressure difference between the interior and exterior of the vesicle. Using Laplace’s law, such a pressure difference is equivalent to a surface tension *σ* = *R*Δ*P/*2, where *R* is the radius of the vesicle.

### B. Mesoscale model of the proteins

In FreeDTS mesoscale model, a protein is modeled as a two-dimensional vector in the plane of a vertex that interacts with the membrane. Here, we focus on proteins whose impact on the membrane is rotationally symmetric within the membrane plane. Therefore, at the mesoscale, their interaction with the membrane is given by

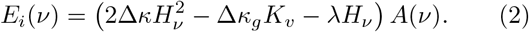

In this equation, Δ*κ* and Δ*κ*_*g*_ are the second-order couplings to the membrane curvature that emerge as a local increase in bending rigidity and Gaussian modulus respectively, representing an increase in the membrane stiffness due to the binding of a rigid protein. *λ* is the first-order coupling parameter to membrane curvature and is related to the local spontaneous curvature *c*_*o*_ induced by the inclusion via *λ* = 2*c*_*o*_(Δ*κ* + *κ*). For more details, see the Supplemental Note 1. Each vertex has an occupancy variable *η*(*ν*), which equals 1 if the vertex is occupied by an inclusion and 0 otherwise.

Therefore, each vertex can host at most one inclusion. 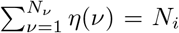 where *N*_*i*_ is the number of inclusions. We define surface coverage *ρ* as the fraction of vertices that are occupied by an inclusion, *ρ* = *N*_*i*_*/N*_*ν*_. For a given configuration of the membrane, the total energy is expressed as

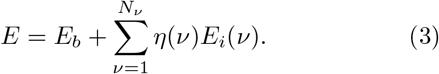

### C. Calibration

To compare the obtained results to biologically relevant systems, we convert the simulation unit length *l*_DTS_ to physical units by assuming that the smallest area associated with a vertex, 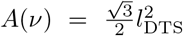, corresponds to the minimal area of a circle with radius *r* enclosing the specific inclusion being modeled (23). This yields 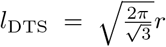. As an example for the B subunit of cholera or Shiga toxin (33, 34) for which their in-plane radius is known to be *r* ≈ 3.5 nm, *l*_DTS_ ≈ 6.7 nm. However, other proteins can be considerably larger. Trimer Piezo1 channel with *r* ≈ 10 − 14 nm (35), would lead to *l*_DTS_ ≈ 18 nm.

### D. System evolution

The system evolution is carried out using a Monte Carlo (MC) scheme with Metropolis algorithm, which samples configurations of the system with Boltzmann’s probability distribution. The dynamically triangulated surface evolves through a series of moves that ensure that all possible triangulations and vertex positions satisfying the imposed constraints are attainable. One MC step consists of *N*_*ν*_ vertex position updates, *N*_*t*_ edge flips (Alexander moves), which switch the mutual connection between two neighboring triangles, and *N*_*i*_ Kawasaki moves, which translate an inclusion from a given vertex to one of its neighboring vertices. For membrane patches under periodic boundary conditions (PBC) in a box, an additional move that changes the box size is required when the system is coupled to a constant frame tension *τ*. In this case, an additional term is added to the energy of the system, *E*_*τ*_ = − *τ A*_*p*_, where *A*_*p*_ is the projected area of the box. The details of the tension-control algorithm can be found elsewhere (23, 39).

### E. System set up

Initialization of the system is performed by randomly placing the inclusions on the surface at the beginning of each run. For each parameter configuration of the system, eight different seeds (which are equivalent to the usage of eight distinct random number sequences) are prepared and run for 5 M MC steps. Every 2000 MC steps, a configuration of the system is stored and used for data analysis. Each data point corresponds to the average of 16000 random configurations coming from eight independent sources. When using the surface-area potential, we fix the compressibility modulus to 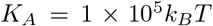, while *A*_*o*_ is obtained by calculating the average surface area of the membrane in absence of constraints or external forces. For a membrane with *τ* = 0, we find an area per vertex of 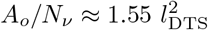.

### F. Clustering measures

To quantify the aggregation of inclusions on the membrane, we employ two complementary clustering measures: the relative average inclusion–inclusion contact and the maximum cluster size. Both measures are based on the concept of nearest neighbors. Two inclusions are defined as nearest neighbors if the vertices they occupy share an edge.

The average number of nearest neighbors per inclusion is defined as

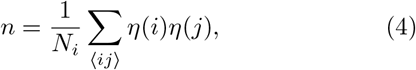

where *N*_*i*_ is the total number of inclusions and the sum runs over all pairs of nearest-neighbor vertices ⟨*ij*⟩, such that each unordered pair of neighboring inclusions is counted twice (once as ⟨*ij*⟩ and once as ⟨*ji* ⟩).

In the absence of direct or indirect interactions between inclusions, they diffuse randomly on the membrane surface. In this case, the expected value of the average number of nearest neighbors is *n*_0_ = *ρm*, where *ρ* is the surface coverage of inclusions and *m* is the average number of edges connected to a given vertex. The difference *n* − *n*_0_ therefore provides a measure of the deviation from random diffusion due to inclusion attraction or repulsion.

To normalize this measure, we divide by the maximum possible positive deviation, *n*_max_ − *n*_0_, where *n*_max_ is the largest value of *n* attainable for a given surface coverage *ρ*. This yields the relative average inclusion–inclusion contact

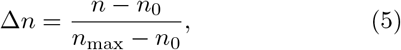

which quantifies local clustering of inclusions. By construction, Δ*n* = 0 corresponds to a random distribution, while Δ*n* = 1 indicates maximal local clustering.

As a complementary, more global measure, we characterize clustering through the maximum cluster size. A cluster is defined as a group of two or more inclusions connected through nearest-neighbor relations, such that any inclusion in the cluster can be reached from any other by successively traversing nearest-neighbor connections. The maximum cluster size, *S*_max_, is defined as the number of inclusions in the largest cluster present on the membrane. We present it normalized by the total number of inclusions, so that it displays the fraction of inclusions present on the system belonging to that cluster. This measure captures large-scale aggregation and is particularly sensitive to the emergence of extended inclusion-rich domains.

## III. RESULTS

### A. Fluctuation-induced forces beat mixing entropy and lead to clustering

First, we investigate whether fluctuation-induced forces can lead to clustering of non-interacting proteins, i.e., in the absence of any direct interactions between the proteins. To this end, we performed DTS simulations of an initially flat membrane patch under PBC containing 1968 vertices. The simulations employed a dynamic box algorithm to maintain fixed frame tension *τ* (39), which was set to zero. The membrane patch is partially covered by proteins that interact only with the membrane and diffuse freely on it.

To isolate fluctuation-induced interactions from curvature-mediated interactions, proteins are modeled as inclusions that locally stiffen the membrane by suppressing bending fluctuations and do not impose any curvature imprint. Thus, we set the inclusion local curvature, *c*_*o*_, to zero and focused on the effects of the inclusion-induced increase in bending rigidity Δ*κ* and Gaussian modulus Δ*κ*_*g*_. A local stability analysis requires 0 ≤ Δ *κ*_*g*_ ≤ 2 (Δ*κ* + *κ*) (see Supplemental Note 2).

We tested various values of inclusion surface coverages (*ρ*) of 5%, 10%, and 20%. For each parameter set, we ran 8 replicas with different seed numbers, each for 5 million MC steps. The simulations were conducted over a wide range of values for Δ*κ* and Δ*κ*_*g*_, while the bare membrane bending rigidity was fixed at *κ* = 20 *k*_*b*_*T*. All shape configurations obtained under these parameters kept the membrane planar, meaning flat on average.

Figure 1(a) shows the relative average inclusion–inclusion contact, Δ*n* = (*n* − *n*_0_)*/*(*n*_*max*_ − *n*_0_), which quantifies local inclusion clustering, as a function of Δ*κ* for several values of the surface coverage *ρ*, with Δ*κ*_*g*_ = Δ*κ*. Here, *n* is the number of nearest-neighbor inclusion pairs, *n*_0_ is the corresponding value for randomly diffusing inclusions at a given surface coverage (i.e., in the absence of any inclusion-induced membrane stiffening), and *n*_*max*_ is the maximum possible value of *n* for a given surface coverage (see Methods for details).

**FIG. 1.**
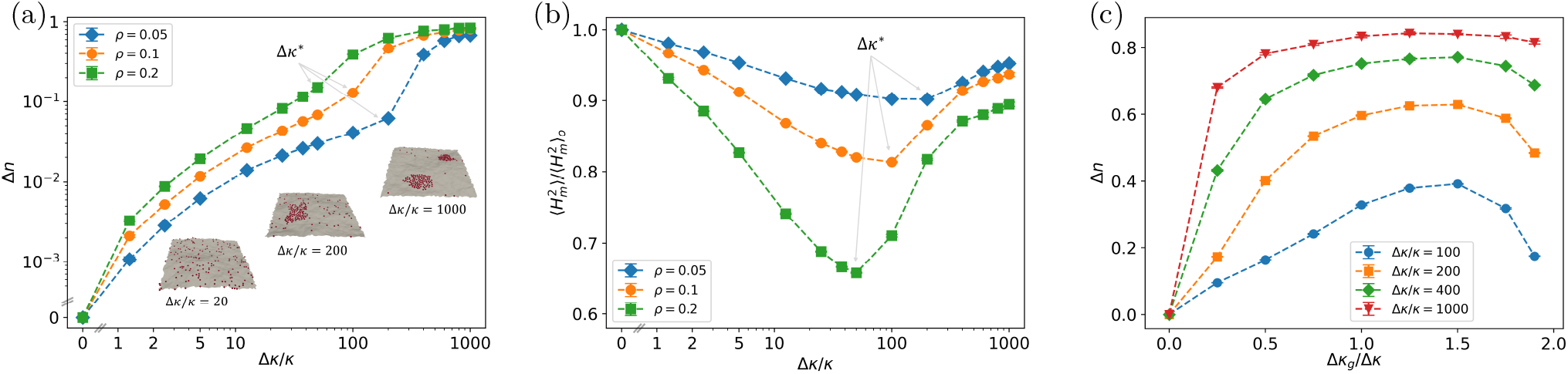
Clustering behavior as a function of inclusion-induced bending rigidity and Gaussian modulus. (a) Average relative inclusion-inclusion contact measure (Δ*n*) as a function of normalized inclusion bending rigidity (Δ*κ/κ*). The inclusion Gaussian modulus is fixed at Δ*κ*_*g*_ = Δ*κ*, and three different surface coverages (*ρ*) are shown. Δ*n* quantifies local inclusion clustering; Δ*n* = 0 corresponds to the random diffusion case, while Δ*n* = 1 corresponds to all inclusions forming a single compact cluster. Surface coverage refers to the fraction of vertices occupied by inclusions. In the simulation snapshots, red spheres mark inclusions (proteins), and gray regions correspond to bare membrane. (b) Average square mean curvature for membrane vertices 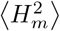 (see main text for a definition) normalized over its value in absence of inclusion stiffness, 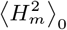. (c) Average relative inclusion-inclusion contact measure (Δ*n*) as a function of normalized inclusion Gaussian modulus (Δ*κ*_*g*_ */*Δ*κ*) for different inclusion bending rigidities Δ*κ/κ*. The surface coverage is fixed at *ρ* = 0.2. Thermodynamic stability requires that 0 ≤ Δ*κ*_*g*_ ≤ 2(Δ*κ* + *κ*).

As Δ*κ* increases from small values (see inset of Fig. 1(a)), Δ*n* increases steadily, indicating enhanced clustering. A crossover from a mild clustered state to an aggregated state is observed as Δ*κ* is increased further, indicated by a sharp rise in Δ*n* (see simulation snapshots of Fig. 1(a), also see Supplemental Movie 1). This crossover is accompanied by the formation of large inclusion clusters containing up to 60–80% of all inclusions (see Fig. S2; see Methods for the definition of cluster).

To characterize this crossover, we evaluate the average square mean curvature for membrane vertices, 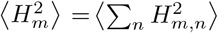, as a function of Δ*κ/κ* (see Fig. 1(b)). Here the sum *n* runs over bare membrane vertices *m*, and ⟨·⟩ denotes an ensemble average. This quantity provides a measure of membrane shape fluctuations and displays a minimum at an inclusion stiffness that we define as crossover bending rigidity Δ*κ*^∗^. In the mild clustering regime (Δ*κ <* Δ*κ*^∗^), increasing inclusion stiffness progressively suppresses membrane fluctuations. In contrast, in the aggregated regime (Δ*κ* > Δ*κ*^∗^), further increases in stiffness lead to enhanced membrane shape fluctuations. Furthermore, Δ*κ*^∗^ coincides with an inflection point of Δ*n*(Δ*κ*) (see Fig. 1(a)). We verified that this behavior is robust with respect to system size (see Fig. S4). Notably, the average energy per vertex remains approximately constant across the entire stiffness range (see Fig. S5). Please note that this is just an observational theory based on the simulated system.

Taken together, these observations suggest that the crossover is governed by a competition between the mixing entropy of the inclusions and the entropy associated with membrane shape fluctuations. In the mild clustering regime, mixing entropy favors a uniform inclusion distribution, at the cost of suppressing membrane shape fluctuations. Once this suppression becomes sufficiently strong, the dispersed state is no longer entropically favorable, and inclusion clustering emerges, driving the crossover to the aggregated regime.

So far, we have shown that fluctuation-induced forces generate interactions between membrane proteins, which in some scenarios alone can lead to clustering. However, to fully observe this phenomenon, long simulations are required, and correct membrane shape fluctuations must be explicitly included in the model. We next ask what simple corrections can be introduced into models where such emergent phenomena are absent by design, for example, in approaches based on optimization procedures or in models that do not include protein stiffness. Such corrections are common in many models: for example, hydrophobic interactions have an entropic origin (7, 49), yet are often represented through effective enthalpic interactions in molecular dynamics and coarse-grained membrane models, including solvent-free approaches and frameworks such as the Martini model (50).

To this end, we perform simulations of a simplified reference system in which inclusions do not interact with the membrane but instead interact directly via a prescribed nearest-neighbor interaction of strength *J*, i.e., *e*_*ij*_ = − *Jδ*_*ij*_. Such a system is equivalent to an Ising model with fixed magnetization or a lattice gas with constant number of particles, for which, in the thermodynamic limit, a continuous phase transition to an ordered state occurs (51, 52). This lattice gas system reproduces the qualitative clustering behavior observed in the membrane-mediated system, including increased clustering with increasing surface coverage (see Fig. S6). The case *J* = 0.5 *k*_*b*_*T* approximates well the leading order interaction for fluctuation-induced forces (26), which is pairwise additive (13). Notably, the clustering measures obtained by the Ising-like system at *J* = 0.5 *k*_*b*_*T* are well below the ones for the real system, especially for high values of inclusion stiffness.

Next, we investigated the effect of Δ*κ*_*g*_ on clustering across the full thermodynamically allowed range. Figure 1(c) shows Δ*n* as a function of Δ*κ*_*g*_ for several values ofΔ*κ* at *ρ* = 0.2. For Δ*κ*_*g*_ = 0, inclusions behave as in the random diffusion case and do not interact with each other. In contrast, in the range 0 < Δ*κ*_*g*_ < 2(Δ*κ* + *κ*), Δ*n* is finite and positive, and displays a maximum located at 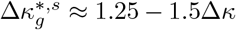. For small Δ*κ*, this maximum is pronounced and broadens as Δ*κ* increases, almost plateauing at very high Δ*κ*. As Δ*κ*_*g*_ approaches the limit 2(Δ*κ* + *κ*) the clustering is severely reduced. These results indicate that clustering requires inclusions to simultaneously suppress both dome-like and saddle-like membrane deformations locally (see Supplemental Note 2). Previous theoretical analysis of two-body interactions (12) have shown that the inclusion-inclusion force exhibits a maximum at a given Gaussian modulus 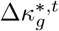 which broadens with increasing Δ*κ* (see Supplemental Note 3). It is interesting that the multi-body behavior follows a similar qualitative behavior.

Overall, these results demonstrate that proteins that bind to the membrane and locally increase its stiffness will cluster. The clustering increases steadily for increasing protein-induced stiffness and for sufficiently stiff proteins, a transition to a fully aggregated state is observed. The protein-induced stiffness required for this transition is on the order of a hundred times that of the membrane. Such high rigidity is unlikely to be common among many membrane proteins. However, for lower rigidity values, relevant to many proteins (53–58), the clustering measures, while mild, are traceable and indicate that fluctuation-mediated forces can contribute to protein clustering. Notably, membrane heterogeneity in biological systems typically manifests as local concentration enrichment rather than complete segregation (59– 63). Please, see the Discussion section for a detailed discussion on this.

The simulated system contained 1968 vertices, corresponding to an average lateral membrane size of approximately 55 *l*_DTS_. Mapping the simulation length unit to physical units therefore depends on the size of the considered proteins. Suppose that the radius of the relevant proteins can vary between 3.5–14 nm; the modeled lateral size then corresponds to membrane patches of sizes ranging from 0.3–1 *µ*m, depending on the protein type (see Methods for details on the calibration procedure). The simulated membrane patch can therefore be viewed as a representative section of a giant unilamellar vesicle (GUV) or a biological membrane with negligible curvature and tension, partially covered by proteins at fixed surface coverage. These conditions are commonly encountered in both biological and experimental settings.

### B. Membrane tension reduces clustering but does not abolish it

Biological membranes are often subject to surface tensions of the order of 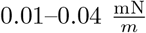, that can reach 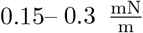 in some cells (64). Tension clearly affects the membrane undulation spectrum (65, 66), and it is reasonable to expect that it also influences fluctuation-induced forces, since these arise from distortions in that spectrum. Indeed, theoretical analyses show that membrane tension weakens such interactions (12). To explore the effects of surface tension, we performed constant-tension simulations with fixed total surface area.

Figure 2 shows the effect of frame tension on inclusion clustering for different inclusion stiffness values, with Δ*κ*_*g*_ = Δ*κ* and surface coverages of *ρ* = 0.05 and *ρ* = 0.2. Clustering is quantified with the inclusion-inclusion contact measure, normalized by its zero-tension value, Δ*n/*Δ*n*(*τ* = 0), which enables direct comparison of the impact of frame tension across different inclusion stiffness values.

**FIG. 2.**
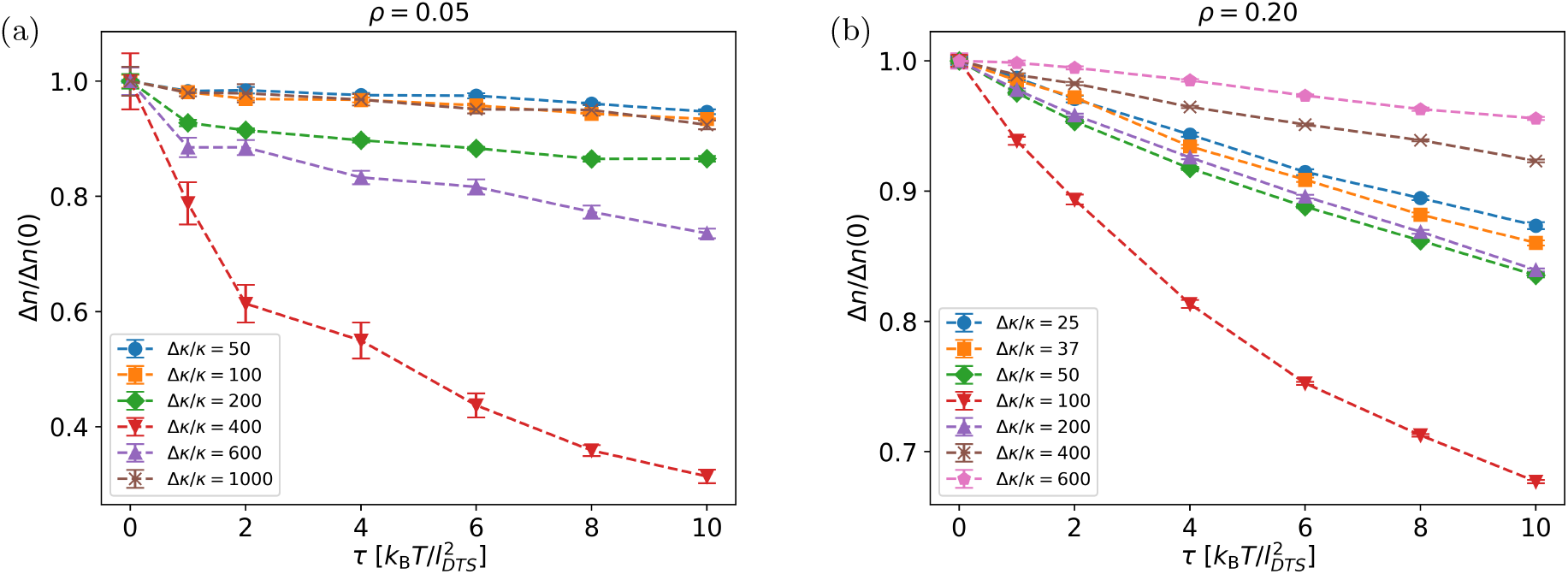
Ensemble average of relative inclusion-inclusion contact, normalized by their corresponding value at zero tension (Δ*n/*Δ*n*(0)) as a function of frame tension (*τ* in units 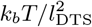) for different inclusion bending rigidities (Δ*κ/κ*) at surface coverages (a) *ρ* = 0.05 and (b) *ρ* = 0.2. The Gaussian modulus is fixed at Δ*κ*_*g*_ = Δ*κ*. The clustering measure Δ*n* is normalized by its zero-tension value Δ*n*(*τ* = 0), allowing comparison of surface tension effects across different inclusion stiffness.

Overall, Δ*n/*Δ*n*(0) decreases as the frame tension increases. The extent of this reduction depends on the difference between the absolute current value of inclusion bending rigidity Δ*κ* and a characteristic value, which we call the variance bending rigidity Δ*κ*^*v*^. This value corresponds to the inclusion stiffness at which the system exhibits the strongest response of clustering to frame tension. The closer Δ*κ* is to Δ*κ*^*v*^, the more sensitive the system is to frame tension, i.e., the more pronounced the reduction in clustering.

For low surface coverage (*ρ* = 0.05), the reduction in clustering at Δ*κ* = Δ*κ*^*v*^ reaches up to approximately 70% at 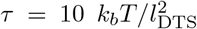, relative to the zero-tension case. In contrast, for higher surface coverage (*ρ* = 0.2), the reduction is smaller, amounting to about 30% under the same conditions. In general, systems at lower surface coverages appear more sensitive to frame tension.

As the difference |Δ*κ* − Δ*κ*^*v*^| increases, the system becomes less susceptible to frame tension. In this regime, the decrease in clustering is remarkably small, being only noticeable for high tensions.

The behavior with respect to the variance bending rigidity Δ*κ*^*v*^ correlates with the variance of the clustering measure Δ*n*, shown in Fig. S10. The variance exhibits a pronounced maximum at Δ*κ* = Δ*κ*^*v*^ and decreases as |Δ*κ* − Δ*κ*^*v*^| increases. The greater sensitivity to frame tension observed for *ρ* = 0.05 compared to *ρ* = 0.2 matches the fact that the variance is also higher for that system.

We emphasize that Δ*κ*^*v*^ is distinct from the crossover bending rigidity Δ*κ*^∗^ defined previously. While Δ*κ*^∗^ marks the point of maximum suppression of membrane shape fluctuations, Δ*κ*^*v*^ identifies the point of maximal clustering fluctuations within the aggregated regime. Consistently, Δ*κ*^*v*^ is located at slightly higher bending rigidity than Δ*κ*^∗^ and corresponds to the onset of strong aggregation accompanied by large clustering fluctuations.

These results can also be understood in terms of the characteristic length of the system, which depends on the bending rigidity and the tension of the membrane 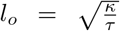. Membrane behavior at distances larger (smaller) than *l*_0_ is dominated by tension (bending) energy. In our simulations, *κ* = 20 *k*_*b*_*T*, giving a crossover length *l*_0_ ranging from 4.47 *l*_DTS_ at the lowest tension 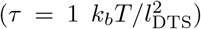 to 1.42 *l*_DTS_ at the highest tension 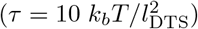, where *l*_DTS_ is the characteristic DTS length scale corresponding to the minimum possible distance between two vertices (or two inclusions). The distance *r* between two inclusions lies within the range of *l*_*o*_, meaning that both surface tension and bending energy dominated fluctuations contribute to the interaction. For *r* > l_*o*_ and inclusion radius *a*, the two-body fluctuation-induced interaction decays as (*a/r*)^8^ and displays a crossover to (*a/r*)^4^ as *r* decreases below *l*_*o*_ (12). As *r* becomes smaller, the interaction strengthens and becomes independent of the surface tension. Relating this to our results, when the difference |Δ*κ* − Δ*κ*^*v*^| is large, the correlation length *ξ* of the interaction remains small. Thus, a decrease in *l*_*o*_ has little effect in the clustering, since *ξ* ≤ *l*_*o*_. In contrast, near the crossover point, *ξ* is large and *l*_*o*_ may become a limiting factor for *ξ*. In that case, decreasing *l*_*o*_ could reduce *ξ*, and consequently the inclusion clustering.

To place the results in a biologically relevant context, we convert *l*_DTS_ to physical units as indicated above (23). The tensions considered here, ranging from 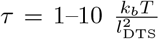, correspond to physical units of approximately 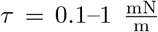 if the protein being modeled as an inclusion is the B subunit of Cholera or Shiga toxin. Our upper limit, 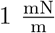, is on the order of the membrane rupture tension (67, 68). Typical biological membrane tensions range from 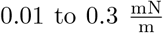, therefore, our simulations span the full range of biologically relevant tensions. With this in mind, we conclude that membrane tension has only a minor effect on fluctuation-induced forces when the system is far from the crossover point Δ*κ*^*v*^. If Δ*κ* = Δ*κ*^*v*^, biologically relevant tensions can reduce clustering by up to 30–40% compared to the tension-free case. In light of previous experimental observations, where clustering of stiff proteins was observed even at high membrane tensions (69), these results might indicate that the experimental setup may have been far from the variance point in the aggregated regime.

### C. Fluctuation-induced forces lead to protein clustering on small vesicles

While flat membranes under PBC are effective models for segments of membranes in, for example, giant unilamellar vesicles (GUVs), where curvature is negligible compared to the protein length scale, many experiments instead use smaller vesicles, such as small unilamellar vesicles (SUVs), where membrane curvature can be important (70–72). For membranes with spherical geometry, it has been shown that membrane curvature affects curvature-mediated forces; however, similar studies have not been conducted for fluctuation-induced forces (30– 32). In contrast with a planar surface under PBC, a vesicle has a nonzero mean and Gaussian curvature, which could impact protein clustering driven by fluctuation-induced forces. To address this gap, we performed simulations of vesicles of different sizes and with varying inclusion surface coverages to test the influence of curvature on fluctuation-induced interactions. Before performing simulations in presence of inclusions, we make sure that the bare simulated vesicle reproduces the expected undulation spectrum for vesicles (see Fig. S1).

Figure 3(a) shows Δ*n* as a function of Δ*κ* for different surface coverages *ρ*, with Δ*κ*_*g*_ = Δ*κ*. Here, Δ*n* follows a similar trend with inclusion stiffness and surface coverage as in the planar case, although the absolute values for Δ*n* are smaller. The transition from the dispersed to the aggregated state is accompanied by local membrane flattening within inclusion clusters, evidenced by a reduction of the average mean and Gaussian curvatures at the inclusion vertices (see Fig. S11). In the aggregated regime, clusters comprise about 5–20% of all inclusions (see Fig. S12) and are evenly distributed across the vesicle surface.

**FIG. 3.**
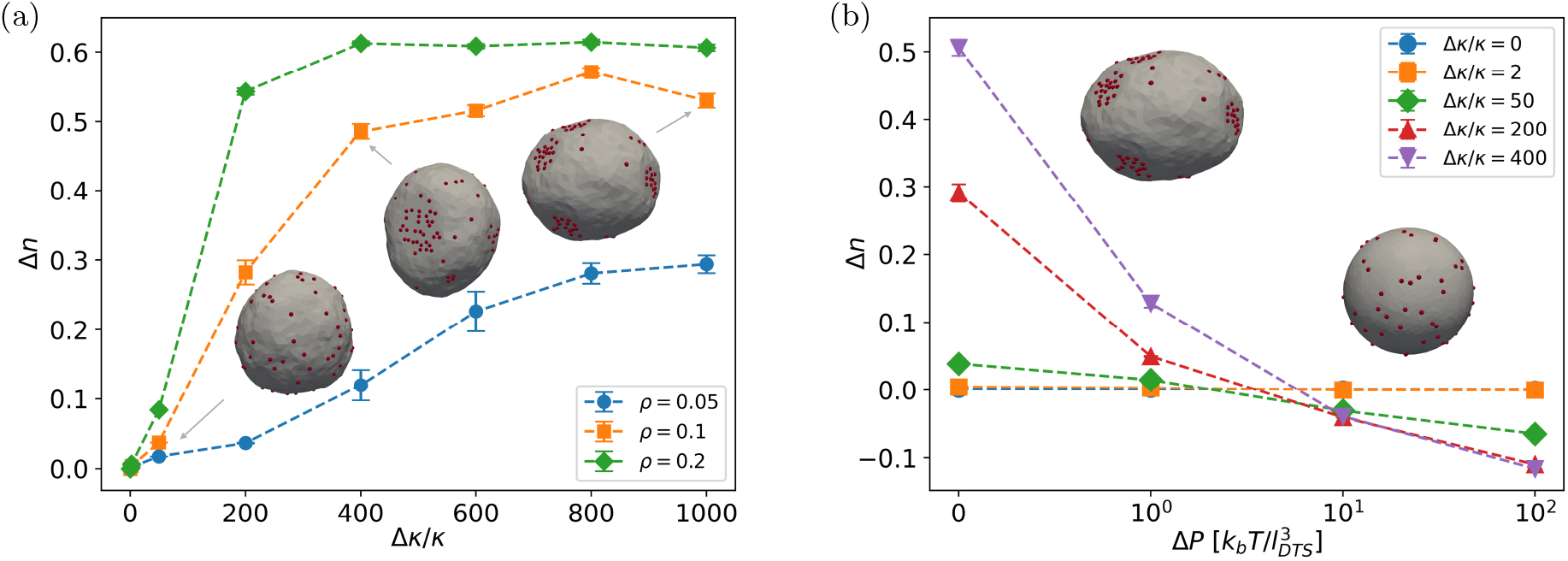
Inclusion clustering on vesicles. (a) Average relative inclusion-inclusion contact measure (Δ*n*) as a function of normalized inclusion bending rigidity (Δ*κ/κ*) for different surface coverages (*ρ*), with the inclusion Gaussian modulus fixed at Δ*κ*_*g*_ = Δ*κ* for spherical vesicle geometry. (b) Average relative inclusion-inclusion contact measure (Δ*n*) as a function of osmotic pressure (Δ*P* in units of 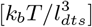) for different normalized inclusion bending rigidities (Δ*κ/κ*).

Next, we include the effect of membrane tension by including a coupling with the vesicle volume of the type − Δ*PV*, where Δ*P* > 0 represents an outward osmotic pressure. The total area of the membrane is kept fixed. At low Δ*P*, increasing Δ*P* hinders the shape fluctuations that facilitate inclusion clustering, resulting in a decrease in Δ*n* (see Fig. 3(b)). Thus, as in the planar case, the addition of an external potential that suppresses membrane fluctuations reduces inclusion clustering. At higher Δ*P*, shape changes are further suppressed, and Δ*n* reaches zero or even negative values, indicating effective repulsion between inclusions. This effective repulsion may originate to keep the volume as high as possible, since inclusions in close proximity could induce a local flattening of the membrane that results in a volume reduction. To examine how vesicle size influences inclusion clustering, we performed simulations with vesicles containing five different number of vertices, *N*_*ν*_ = 650, 802, 1252, 1570, 1986. The results are displayed in Fig. S13, where Δ*n* is plotted against Δ*P* for different vesicle sizes and different inclusion stiffness. Across all inclusion stiffness and osmotic pressure, clustering increases with vesicle size. Several factors may contribute to this trend. First, larger vesicles exhibit stronger shape fluctuations (73), which implies that fluctuation-mediated forces are enhanced compared to smaller vesicles. Second, forming locally flat membrane regions is likely easier in larger vesicles, since their average mean curvature decreases with size. In other words, membrane flattening is easier when the initial curvature is lower. Both effects favor inclusion clustering in larger vesicles and thus, shape fluctuations and curvature constraints are the main drivers of the observed vesicle-size dependence.

Taken together, these results show that stiff proteins can indeed cluster on the surface of SUVs at sufficiently low osmotic pressure. Note that for these simulations we fix the average area per vertex as we change *N*_*ν*_, and thus increasing number of vertices is equivalent to increasing the area of the simulated surface. Thus, the radius of the simulated vesicle is 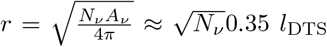. The smaller and bigger size of simulated vesicle are equivalent to 9 and 16 *l*_DTS_ of the simulated vesicle. If we consider that the proteins modeled in these simulations are Shiga toxins, the simulated vesicle radii range around *R* ≈ 60– 120 nm, corresponding to the typical size of SUVs. The osmotic pressure values can also be converted from simulation to physical units, giving 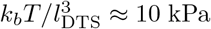. Typical osmotic pressures experienced by liposomes are in the kPa range: for example, lysis thresholds in aspiration experiments fall between 3–6 kPa (74) and FRET-based liposomal sensors operate within 0–300 kPa (75). Overall, this indicates that under biologically relevant osmotic pressures, the inclusions will be able to cluster. The clustering might be quite sensitive to an increase in osmotic pressure when compared to an increase of the same magnitude in frame tension in the planar surface case. Protein repulsion could be observed for very high osmotic pressure close to the membrane rupture.

### D. Rigid proteins absorb less rigid proteins through fluctuation-induced forces

In biological membranes, it is common for proteins of different species to cluster together to carry out specific functions (76, 77). These proteins interact with the membrane in different ways and with varying strengths, which we represent in our model by assigning them distinct model parameter values. It is therefore of great interest to investigate how long-range fluctuation-induced forces may contribute to their clustering. To explore complex systems beyond single-protein-type scenarios, we performed simulations of membranes containing two types of inclusions, each characterized by distinct values 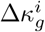 and Δ*κ*^*i*^, where *i* = 0, 1 denotes the inclusion type. Both inclusions are present at the same concentration *ρ*_0_ = *ρ*_1_ = 0.05.

Figure 4 shows the relative average inclusion-inclusion contact measure for inclusions of the same type, *n*_*ii*_, and for inclusions of different type, *n*_*ij*_, plotted as a function of the Gaussian modulus difference 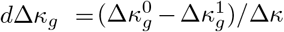 for 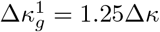 and varying 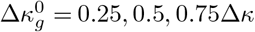. The inclusion bending rigidity is kept fixed, Δ*κ*^0^ = Δ*κ*^1^ = Δ*κ*. In addition to interactions between inclusions of the same type, interactions between inclusions of different type also take place. Our results show that the clustering follows *n*_11_ > *n*_01_ > *n*_00_. In previous sections, we showed that inclusions of type 1 exhibit higher measures of clustering than inclusions of type 0 in isolation (see Figure 1). This explains why *n*_11_ > *n*_00_. However, the observation that *n*_01_ > *n*_00_ is a new and unintuitive feature that is relevant for complex systems composed of more than one type of protein. We study other combinations of 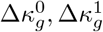 and Δ*κ*^0^, Δ*κ*^1^ observe that, in general, if one inclusion type displays lower measures of clustering in isolation than the other, it is more strongly attracted to the other inclusion type than to itself (see Fig. S14). In other words, less stiff proteins are more strongly attracted to stiffer proteins than to themselves.

**FIG. 4.**
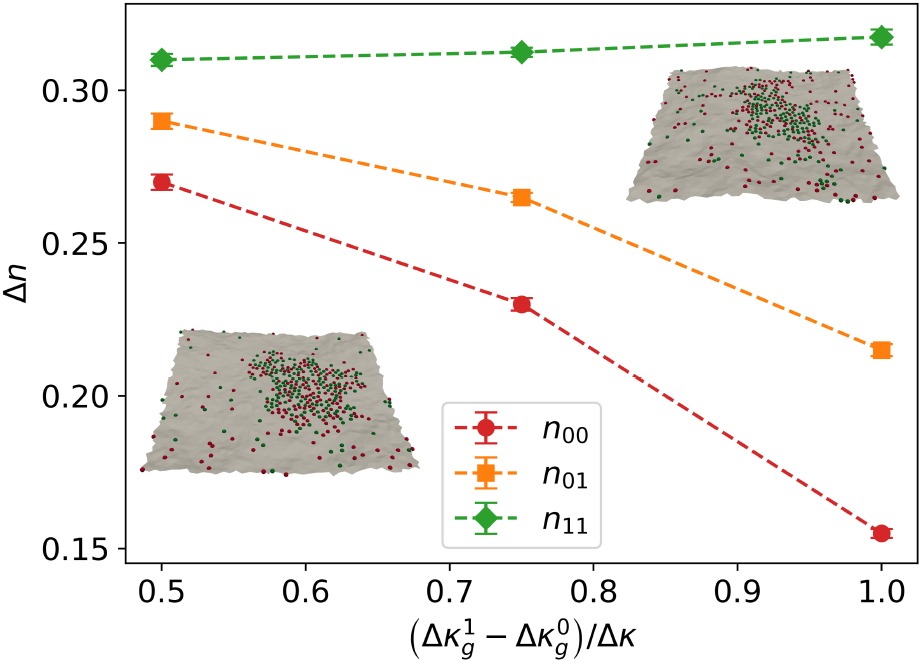
Average relative inclusion-inclusion contact measure (Δ*n*) as a function of the normalized difference in Gaussian modulus, 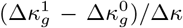, between two inclusion types. Here, *n*_00_ corresponds to Δ*n* for inclusions of type 0 (with 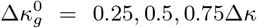 shown in red in the simulation frames), *n*_11_ corresponds to Δ*n* for inclusions of type 1 (with 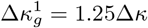 shown in green in the simulation frames). *n*_01_ corresponds to Δ*n* between inclusions of type 0 and type 1. The inclusion bending rigidity is Δ*κ*^0^ = Δ*κ*^1^ = 500*κ*, and the surface coverage is *ρ*_0_ = *ρ*_1_ = 0.05.

These results indicate that stiffer proteins can act as nucleation centers, promoting the clustering of less stiff proteins.

### E. Curvature imprint can reduce clustering but proteins find their way

So far, we have considered proteins that locally increase the stiffness of the lipid bilayer. However, a common characteristic of many membrane-associated proteins is their ability to induce local membrane curvature. This can occur through peripheral binding to the membrane surface, through transmembrane proteins with conical shapes, or via specific interactions with the lipid environment. In such cases, the protein imposes a preferred local curvature on the membrane (8, 33, 34, 78, 79). To model curvature-inducing, stiff proteins, we introduce a nonzero local curvature *c*_*o*_ in the inclusion parameters (2). The mean curvature imprinted by an inclusion is obtained by minimizing the local energy of a vertex containing it, yielding: 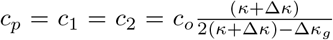. We performed simulations for different values of Δ*κ/κ* with Δ*κ*_*g*_ = Δ*κ*, and for 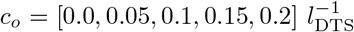. A low surface coverage (*ρ* = 0.1) is chosen to isolate the effects of individual inclusions and avoid membrane shape remodeling driven by curvature instabilities that arise at high inclusion surface coverages or regions of strong local curvature (39, 80, 81). Membrane bending rigidity is set to *κ* = 10 *k*_*b*_*T* for improved sampling.

We first examine the effects of membrane-mediated interactions in the low-deformation regime. To do so, an external frame tension 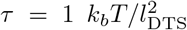 is applied (the surface area *A* is kept constant). In the low stiffness regime, the clustering depends widely on the inclusion local curvature (see inset of Fig. 5(a)). For 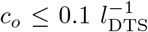 increasing stiffness consistently enhances aggregation. By contrast, an effective repulsion indicated by Δ*n <* 0 is observed for 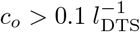. This decrease in inclusion clustering is followed by an increase as the inclusion stiffness increases further. Previous studies have shown that curvature-mediated forces between rigid inclusions are primarily repulsive, especially for small curvature imprints. This repulsion scales with the square of the curvature imprint, 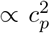 (9, 13). Multi-body effects and tension have been shown to modulate clustering effects (82–84). Our results agree well with this picture: at sufficiently high *c*_*o*_, curvature-mediated repulsion dominates over fluctuation-induced attraction in the low stiffness regime. As the stiffness is increased further, the curvature imprint plateaus, leaving only the fluctuation-induced components to grow, which restores clustering.

**FIG. 5.**
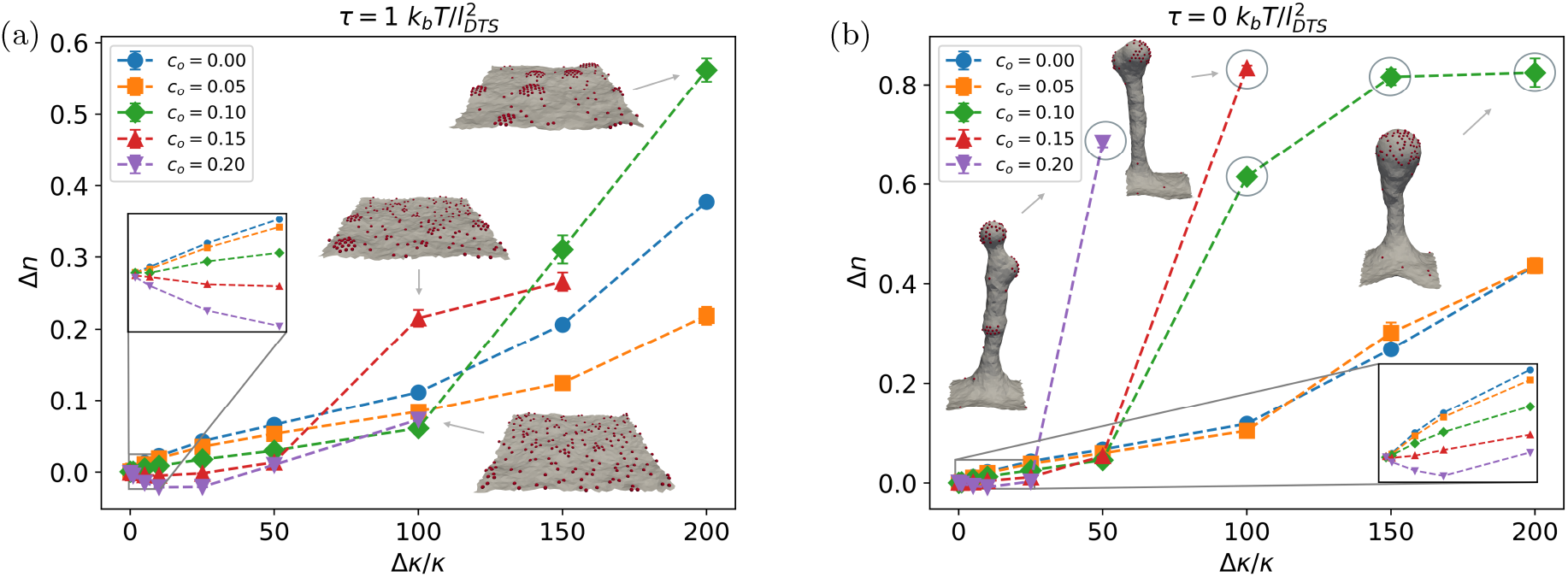
Average relative inclusion-inclusion contact measure (Δ*n*) as a function of normalized bending rigidity (Δ*κ/κ*) for inclusions inducing five different local curvatures co (in units of 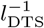), shown for frame tensions (a) 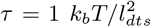 and (b) 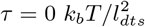. Circled points correspond to points that were run further 30 *M* steps.

For very stiff inclusions and sufficiently high local curvature, 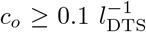, a sudden increase in clustering is accompanied by a rise in total average mean curvature (see Fig. S15). This suggests that even in presence of a mild tension, inclusions can collectively deform the membrane into spherical caps that then act as clustering centers (see simulation frames in Fig. 5(a)). The resulting spherical caps resemble those also obtained in previous studies for curvature-inducing, stiff nanoparticles, suggesting that we recover their observed phenomena as a regime in our parameter space (26, 27). In this regime, aggregation driven by fluctuation-induced forces is further enhanced by attractive, curvature-mediated interactions that emerge when inclusions collectively generate local membrane deformations.

To further probe this effect, we ran simulations at zero frame tension 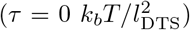. The results follow the same general trends observed at 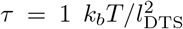 (seeFig. 5(b)), but the transition from planar to deformed state occurs at lower inclusion stiffness. For simulations where the membrane undergoes shape remodeling departing from a flat configuration, the simulations have been run further 30 *M* steps. The resulting membrane deformations are much stronger; the membrane can undergo shape remodeling and membrane invaginations are formed (see simulation frames in Fig. 5(b) and Fig. S15). The inclusion parameters determine the shape of the membrane and inclusion distribution of the final configurations, which resemble membrane tubes and buds. Prior to the formation of the tubes, the inclusions form clusters with spherical cap, which then merge and induce tubulation and budding (see Fig. S16). Previous studies that observed the formation of buds for similar nanoparticles as the ones used here required the use of external forces to induce the formation of buds (29). Here, the buds and tubes emerge spontaneously. Importantly, the membrane shape remodeling observed here differs from budding driven by membrane softening mechanism (39), which requires higher entropy of mixing (e.g higher surface coverage) or larger curvature imprints. To ensure this, we provide measures of the average square mean cur vature for membrane vertices 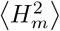 for inclusion parameters that do not give rise to membrane shape remodeling and confirm that indeed, no membrane softening occurs prior to the transition (see Fig. S17). Note that inclusions generating extremely large curvatures can experience attractive curvature-mediated interactions, potentially leading to clustering (85). However, this curvature regime is far beyond the curvatures typically induced by membrane proteins (33, 34, 39) and is instead more relevant for colloidal particles (14). These curvature-induced interactions have been invoked to explain membrane budding induced by nanoparticles, even in cases where the induced curvature lies below the threshold required for an attractive two-body interaction (86). Reinterpreting the budding observed in reference (86) in light of our results, one may speculate that the interplay between fluctuation-induced forces and curvature-mediated interactions produces a nontrivial effective interaction that drives clustering and subsequent bud formation.

In summary, while curvature imprint can produce an effective repulsion between stiff proteins, it can also enhance clustering once fluctuation-induced attraction becomes dominant. This reveals a nontrivial interplay: when curvature-mediated repulsion is too weak to suppress clustering, collective curvature generation instead amplifies aggregation. This aggregation can even drive large-scale membrane remodeling, highlighting a synergistic mechanism for clustering and budding.

Finally, the curvature imprints used here 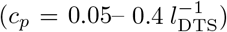 correspond to *c*_*p*_ = 0.008–0.06 nm^−1^ in physical units for proteins such as Shiga or cholera toxin. These values fall within the experimentally measured range ~ 0.03 nm^−1^ (33, 34).

## IV. DISCUSSION

In this work, we investigated the lateral organization of non-interacting membrane inclusions that locally modify membrane stiffness and/or induce membrane curvature across a broad range of mesoscale parameters and inclusion concentrations. The model captures key features of both peripheral proteins tightly bound to the membrane and transmembrane proteins, while enabling continuous quantification of spatial clustering across the full parameter regime. Whereas previous computational studies explored membrane-mediated interactions only in restricted parameter regimes (26–29), our approach provides a systematic view of how clustering emerges across a wide range of protein mechanical properties and membrane conditions.

Our results show that fluctuation-mediated interactions progressively drive proteins away from random mixing as protein rigidity or surface coverage increases, leading from weak clustering to complete segregation at sufficiently high protein rigidity. Unlike previous computational works, which were limited to identifying the existence of such a transition (24), our approach reveals that significant clustering already emerges well before complete segregation is reached. This indicates that fluctuation-mediated interactions can significantly reorganize proteins even in regimes where macroscopic phase separation is absent.

A key determinant of this clustering is the local Gaussian modulus induced by the proteins. Proteins must suppress both saddle and dome membrane fluctuations to cluster efficiently, with clustering consistently peaking at Δ*κ*_*g*_ ≈ 1.25 − 1.5Δ*κ*. While the importance of Gaussian rigidity has been established in the asymptotic two-body limit (9, 12), its role in many-body clustering had not been systematically explored.

Despite the large literature devoted to pairwise asymptotic expressions for fluctuation-induced interactions, our results demonstrate that such approximations substantially underestimate clustering, even in the limit of infinitely rigid proteins. Previous analytical works demonstrated that properly accounting for higher-order terms can strengthen the two-body interaction by roughly one or two orders of magnitude for infinitely rigid inclusions (12, 13, 87). Including non-additive multi-body effects on top of this enhancement could further explain the high clustering measures we observe for very stiff proteins.

Beyond inducing stiffness, many proteins induce a preferred spontaneous curvature (8, 33, 34, 57, 78, 79). We find that curvature-mediated interactions can produce an effective repulsion when they outweigh fluctuation-induced attractions, consistent with previous theoretical works (9, 13, 82–84). However, in an intermediate regime, fluctuation-mediated attraction overcomes this repulsion, leading to both clustering and spontaneous membrane tubulation without requiring the use of external forces, in contrast with previous computational works (29). Notably, tubulation occurs at lower protein rigidity than that required for significant clustering in flat membranes, revealing a synergy between curvature-mediated and fluctuation-induced forces that neither mechanism produces independently.

We further explored several physical regimes that have received limited or no attention in the context of multi-body fluctuation-induced forces, including membrane tension, closed vesicle geometries, and mixed protein systems.

The impact of membrane tension on fluctuation-mediated forces has been explored in a limited number of theoretical studies (12, 13, 87), restricted to asymptotic expressions in tension-dominated regimes, where tension is predicted to strongly suppress pairwise forces, changing the interaction decay from 1*/r*^4^ to 1*/r*^8^. Here, we go beyond this limit and demonstrate that, despite this strong effect at the two-body level, surface tension reduces clustering only modestly in the multi-body case, with pronounced effects appearing exclusively near the crossover bending rigidity.

We also examined fluctuation-mediated interactions on closed vesicles, where the membrane possesses intrinsic curvature, as in synthetic vesicle systems such as SUVs. At sufficiently low osmotic pressure difference, proteins cluster similarly to the planar case, although clusters remain smaller and more evenly distributed across the surface. This indicates that fluctuation-mediated forces in vesicles may generate small, localized protein clusters with functional relevance, such as in signaling (88).

In binary mixtures of proteins with different rigidities, stiffer proteins act as nucleation centers for softer proteins, promoting heterogeneous cluster formation. This suggests that fluctuation-mediated interactions may contribute to the assembly of mixed protein complexes and reinforce lateral membrane heterogeneity in biologically relevant multicomponent systems (76, 77).

One should note that these results have been obtained using a dynamically triangulated surface representation of the membrane together with a single particle-based protein model, and therefore do not accurately capture the close-proximity regime, i.e., separations at which the lateral shape and detailed structure of the inclusions become important. Analytical studies have shown that the interaction scales as 1*/d* in both the bending and tension-dominated regimes, albeit with different prefactors, where *d* denotes the edge-to-edge separation between the inclusions (8, 11). Consequently, the interaction energy diverges as *d →* 0, implying that the energetic gain upon aggregation becomes highly sensitive to the microscopic cutoff length below which the continuum description, together with the associated membrane undulation modes (89, 90), loses physical validity. At this scale, other degrees of freedom and their associated fluctuations, such as membrane thickness and lipid tilt, become important. To study short-range fluctuation-mediated interactions, methods such as molecular dynamics and dissipative particle dynamics are more appropriate (8, 33). Moreover, in our model, the microscopic cutoff is set by the simulation unit length *l*_DTS_, which we convert to physical units by setting it equal to the size of the protein under consideration. While this is a good approximation for many proteins, whose characteristic size is on the order of the membrane thickness, the protein size and the cutoff length are, in general, independent. For proteins smaller than the membrane thickness, however, this identification is no longer appropriate, as the additional degrees of freedom that become relevant at that scale are not captured by our model.

Our results further indicate that complete segregation requires protein-induced rigidities more than two orders of magnitude larger than the surrounding membrane, suggesting that this regime is likely restricted to specialized systems with exceptionally strong membrane coupling. Reports of near-complete protein segregation or very high clustering measures remain relatively uncommon in the experimental and computational literature, where protein assemblies more typically consist of small dynamic clusters rather than macroscopic segregated domains (59–63). Thus, the biologically more relevant regime likely corresponds to proteins that stiffen the membrane by only a few to ten or, maximum, hundred fold. In this regime, fluctuation-mediated interactions do not fully dominate mixing entropy but instead act as tunable, cooperative forces that can promote clustering. In synergy with other weak interactions, such as depletion forces or transient direct contacts (91, 92), they may therefore contribute significantly to protein organization without necessarily inducing macroscopic phase separation in the cellular context. Importantly, proteins that also impose local spontaneous curvature may access strong fluctuation-mediated organization at substantially lower rigidity, as demonstrated by our curvature results above.

Current estimates of protein-induced membrane stiffening remain scarce and span a wide range. While isolated proteins are typically two to three orders of magnitude stiffer than lipid membranes based on Young modulus measurements (93, 94), these values do not directly translate into the effective stiffness induced on the membrane, which also depends on protein geometry, binding mode, and lipid environment (5). In general, proteins that bind tightly to the membrane over a large lateral area, as well as transmembrane proteins with large lateral area, are expected to induce substantial local stiffening. Examples of these proteins would be Cholera or Shiga toxin (33, 34), Annexins (95, 96) or Piezo1 (35), for which no such measurements exist yet. In fact, high measures of Shiga toxin clustering on the membrane depended on how tightly the protein was bound to the membrane, suggesting that fluctuation-mediated forces are the only plausible theory capable of explaining its almost complete clustering (8). Available membrane-coupled estimates nonetheless suggest a broad spectrum of induced rigidities: molecular dynamics studies report that *β*-barrel transmembrane proteins can locally stiffen membranes by approximately 90–190 fold (97); clathrin coats increase membrane rigidity by up to roughly 20-fold (98, 99), whereas indirect measurements for the rigidity of other proteins appear to indicate more moderate rigidity changes, on the order of two to tenfold relative to the bare membrane (53–58). Together, these estimates suggest that many membrane-associated proteins may operate in a regime where fluctuation-mediated interactions are sufficiently strong to promote local clustering or lateral reorganization, but not strong enough to drive complete segregation.

As we showed in Fig. 1(b), the clustering reported here is entropic in origin. Entropy-driven ordering is common in biological systems. For example, depletion interactions between membrane proteins arise from the entropy of the mediating medium, specifically the translational and configurational entropy of lipids (5, 7, 91, 92). In contrast to lipid density correlations, which are short-ranged, membrane fluctuations are long-ranged, which makes these effects particularly interesting, as they can operate over relatively larger distances and are not strongly dependent on membrane composition, in particular on lipid chain length (100).

An important future direction is the development of bottom-up methods to systematically determine the three mesoscale parameters used in this work: the local bending rigidity Δ*κ*, Gaussian modulus Δ*κ*_*g*_, and spontaneous curvature *c*_*o*_. While molecular dynamics studies have successfully quantified protein-induced spontaneous curvature from atomistic trajectories (33, 34, 57), systematic protocols for extracting local elastic moduli remain largely undeveloped. In the longer term, combining molecular simulations with machine learning or structure prediction tools such as AlphaFold may enable direct estimation of mesoscale membrane mechanical parameters from protein sequence or structure (101). Once such parameters are available, the clustering maps generated here provide an immediate framework for predicting the extent of fluctuation-mediated protein organization in specific membrane systems.

## V. CONCLUSIONS

In conclusion, our study demonstrates that long-range fluctuation-mediated forces can promote protein aggregation when proteins significantly increase the local stiffness of the membrane. Remarkably, when proteins also induce spontaneous curvature, the stiffness threshold required for aggregation is reduced, and clustering is accompanied by pronounced membrane deformations. These deformations can ultimately drive membrane remodeling and budding, revealing a synergistic mechanism by which stiffness and curvature co-regulate protein organization and membrane shape. Furthermore, these findings suggest that protein accumulations observed in experimental membrane imaging data may result from (or be enhanced by) membrane fluctuations rather than direct affinities, highlighting the need for careful consideration when interpreting membrane-imaging data.

## DATA AVAILABILITY STATEMENT

Source data for the graphs presented in this work are available at https://doi.org/10.5281/zenodo.20365837. The simulation schemes, tutorials and analysis code for data analysis are publicly available at https://github.com/weria-pezeshkian/FreeDTS/wiki/Multi-body-Fluctuation-Induced-Forces.

## AUTHOR CONTRIBUTION

W.P. conceived the original idea. A. B. V. performed simulations and analyzed the results. A. B. V. and W.P. wrote the manuscript. Both authors discussed the results and commented on the manuscript at all stages.

## DECLARATION OF INTERESTS

The authors declare no competing interests.

## ACKNOWLEDGMENTS

We thank M. Wood for comments and feedback on this manuscript. This research is supported by the Novo Nordisk Foundation (grant No. NNF18SA0035142 and NNF22OC0079182) and Independent Research Fund Denmark (grant No. 10.46540/2064-00032B). The Tycho supercomputer hosted at the SCIENCE HPC center at the University of Copenhagen was used for supporting this work.

## Notes

### Competing Interest Statement

The authors have declared no competing interest.

### Summary of Updates

We upload the final version of the manuscript with a small addition to the discussion and a correction of the normalization factor in equation 4.

