## Supporting information for "Multi-body Fluctuation-Induced Forces Between Membrane Proteins: Insights from Mesoscale Simulations"

for

### Supplemental Notes

### 1 Symmetric Inclusion Model

By the Fundamental Theorem of Surface Theory, a surface is uniquely determined, up to rigid motions (translations and rotations), by its metric tensor and curvature tensor (Shape Operator). Consequently, the elastic free energy of a membrane patch, whether or not it contains inclusions, must depend only on geometric invariants of the surface.

For a fluid membrane, the relevant local geometric invariants are the mean curvature $H$, the Gaussian curvature $K$, and the area element $dA$. Since the membrane energy is extensive in the number of constituent molecules (e.g., lipids), the total energy must be additive in area and therefore expressible as a surface integral over a local energy density.

Assuming that the radii of curvature are much larger than the membrane thickness $h$, the energy density may be expanded systematically in powers of curvature. Retaining terms up to quadratic order yields the most general local bending energy consistent with rotational invariance:

$$E_B = \int \left(a_0 + a_1 H + a_2 H^2 + a_3 K\right) dA, \tag{S1}$$

where the coefficients $a_i$ depend on the membrane composition and on the presence of inclusions. Higher-order curvature terms are in principle allowed, but they are suppressed by powers of the small parameter $h/R$, where $R$ denotes a characteristic radius of curvature. Thus, for $h/R \ll 1$, the quadratic expansion provides the dominant contribution to the membrane elastic energy. The constant term $a_0$ is related to the chemical potential. We ignore it from the rest of our calculation by assuming that we are dealing with systems with constant total area.

For a uniform system, Eq. (S1) reduces to the Helfrich Hamiltonian upon identifying

$$a_1 = -2\kappa\bar{C}, \qquad a_2 = 2\kappa, \qquad a_3 = -\kappa_g. \tag{S2}$$

The bending energy then takes the form

$$E_B = \int \left[\frac{\kappa}{2}(2H - \bar{C})^2 - \kappa_g K\right] dA. \tag{S3}$$

Here, $\kappa$ and $\kappa_g$ are the bending rigidity and Gaussian modulus, respectively, while $\bar{C}$ denotes the spontaneous curvature. For transbilayer-symmetric membranes, $\bar{C} = 0$. We adopt the sign convention in which the Gaussian curvature term appears with a minus sign, so that positive $\kappa_g$ corresponds to an energetically unfavorable Gaussian curvature contribution. Note that some literature uses the opposite sign convention.

For a dynamically triangulated membrane, the surface is discretized into vertices $\nu$, and the continuum energy is replaced by a discrete sum over local surface elements:

$$E_B = \sum_{\nu=1}^{N_\nu} \left[-\lambda(\nu)H(\nu) + \frac{\kappa(\nu)}{2}\big(2H(\nu)\big)^2 - \kappa_g(\nu)K(\nu)\right] A(\nu), \tag{S4}$$

where $A(\nu)$ is the area associated with vertex $\nu$, and $\lambda(\nu)$, $\kappa(\nu)$, and $\kappa_g(\nu)$ are the local elastic parameters.

We now consider a membrane containing two types of vertices: ordinary membrane vertices and vertices occupied by inclusions. Let

$$\eta(\nu) = \begin{cases} 1, & \text{if an inclusion is present at vertex } \nu, \\ 0, & \text{otherwise.} \end{cases} \tag{S5}$$

Assuming that the bare membrane is symmetric, the spontaneous curvature vanishes in the absence of inclusions. By considering that the inclusions are symmetric under in-planer rotations, the local elastic moduli are then written as

$$\kappa(\nu) = \kappa_o + \eta(\nu)\Delta\kappa, \tag{S6}$$

$$\kappa_g(\nu) = \kappa_{go} + \eta(\nu)\Delta\kappa_g, \tag{S7}$$

$$\lambda(\nu) = \eta(\nu)\lambda. \tag{S8}$$

Substituting these expressions into Eq. (S4) gives

$$E_B = \sum_{\nu=1}^{N_\nu} \left[ \frac{\kappa_o}{2} (2H(\nu))^2 - \kappa_{go} K(\nu) \right] A(\nu) \\ + \sum_{\nu=1}^{N_\nu} \eta(\nu) \left[ -\lambda H(\nu) + \frac{\Delta\kappa}{2} (2H(\nu))^2 - \Delta\kappa_g K(\nu) \right] A(\nu). \tag{S9}$$

The first term corresponds to the bending energy of the symmetric membrane, while the second term represents the modification induced by inclusions.

Completing the square in the mean-curvature contribution allows the energy to be rewritten in a form that explicitly separates membrane vertices with and without inclusions:

$$E_B = \sum_{\nu=1}^{N_\nu} (1 - \eta(\nu)) \left[ \frac{\kappa_o}{2} \big(2H(\nu)\big)^2 - \kappa_{g0} K(\nu) \right] A(\nu) \\ + \sum_{\nu=1}^{N_\nu} \eta(\nu) \left[ \frac{\kappa_o + \Delta\kappa}{2} \big(2H(\nu) - c_0\big)^2 - (\kappa_{go} + \Delta\kappa_g) K(\nu) \right] A(\nu), \tag{S10}$$

where the spontaneous curvature induced by the inclusion is defined through

$$\lambda = 2(\kappa_o + \Delta\kappa) c_o. \tag{S11}$$

Thus, vertices without inclusions obey the standard symmetric Helfrich Hamiltonian with zero spontaneous curvature, while vertices containing inclusions acquire a local spontaneous curvature $c_o$ together with modified elastic moduli.

### 2 Thermodynamic stability and intuition for inclusion Gaussian modulus

In the following, we demonstrate that the increase in Gaussian modulus by protein, $\Delta\kappa_g$, introduced in the main text is constrained by stability considerations to the range $0 \le \Delta\kappa_g \le 2(\Delta\kappa + \kappa)$. We also provide physical intuition for the role of $\Delta\kappa_g$ by discussing how its variation corresponds to moving between the two extremes of the thermodynamically stable region. The bending energy of a vertex containing an inclusion is given by the following expression:

$$E(c_1, c_2) = A \left( \frac{\kappa + \Delta\kappa}{2} (c_1^2 + c_2^2) + (\Delta\kappa + \kappa - \Delta\kappa_g) c_1 c_2 \right) \tag{S12}$$

where $A$ and $c_1, c_2$ are the area and principal curvatures associated with the given vertex. This energy can be rewritten as

$$E(c_1, c_2) = \frac{A}{2} \begin{pmatrix} c_1 & c_2 \end{pmatrix} \mathbf{M} \begin{pmatrix} c_1 \\ c_2 \end{pmatrix}, \tag{S13}$$

where the matrix $\mathbf{M}$ is

$$\mathbf{M} = \begin{pmatrix} \kappa + \Delta\kappa & \kappa + \Delta\kappa - \Delta\kappa_g \\ \kappa + \Delta\kappa - \Delta\kappa_g & \kappa + \Delta\kappa \end{pmatrix}. \tag{S14}$$

By diagonalizing this matrix and determining its eigenvectors, we obtain the following expression for the energy:

$$E = \frac{A}{4} \left[ (2(\kappa + \Delta\kappa) - \Delta\kappa_g) (c_1 + c_2)^2 + \Delta\kappa_g (c_1 - c_2)^2 \right]. \tag{S15}$$

For the membrane segment to be strictly stable against arbitrary curvature deformations, both eigenvalues of the matrix $\mathbf{M}$ must be strictly positive. This requirement leads to the conditions $2(\Delta\kappa + \kappa) \ge \Delta\kappa_g$, and $\Delta\kappa_g \ge 0$, which together imply $0 \le \Delta\kappa_g \le 2(\Delta\kappa + \kappa)$. Furthermore, note that while the first term penalizes the formation of domes (shapes that obey $c_1 c_2 > 0$), the second term penalizes the formation of saddle shapes $c_1 c_2 < 0$. Thus, $\Delta\kappa_g = 0$ corresponds to the case in which saddle shapes are not penalized energetically, while the other extreme, $\Delta\kappa_g = 2(\Delta\kappa + \kappa)$, corresponds to the case in which domes are not penalized energetically.

### 3 Theoretical prediction for thermal Casimir-like force

Following Lin et al. [1], the thermal Casimir-like force between two disk inclusions of radius $a$ in the asymptotic limit can be expressed as

$$\beta \mathcal{F}_C = -A \left(\frac{a}{r}\right)^n, \quad \beta \equiv \frac{1}{k_B T}, \quad r \gg a. \tag{S16}$$

For membranes with zero surface tension, the exponent takes the value $n = 4$. If the inclusions locally increase the bending rigidity by $\Delta\kappa$ and the Gaussian modulus by $\Delta\kappa_g$, the prefactor $A$ is given by

$$A = \left[\frac{\Delta\kappa_g}{4\kappa + \Delta\kappa_g}\right] \left[\frac{4\Delta\kappa_g}{4\kappa + \Delta\kappa_g} + \frac{4\Delta\kappa - 2\Delta\kappa_g}{2(\kappa + \Delta\kappa) - \Delta\kappa_g}\right]. \tag{S17}$$

In the rigid-disk like limit ($\Delta\kappa, \Delta\kappa_g \to \infty$), one recovers $A = 6$, in agreement with previous studies [2]. Thermodynamic constraints impose a limit for the local increase in Gaussian modulus to $0 \geq \Delta\kappa_g \geq 2(\kappa + \Delta\kappa)$. The prefactor $A$ as a function of the normalized inclusion Gaussian modulus $\Delta\kappa_g/\Delta\kappa$ is plotted in Fig. S7 for different values of the inclusion bending rigidity $\Delta\kappa/\kappa$. It displays a maximum at a given value of the inclusion Gaussian modulus $\Delta\kappa_g^{t,*} = \Delta\kappa_g^{t,*}(\Delta\kappa)$ that depends on the inclusion bending rigidity. As $\Delta\kappa$ increases, the maximum broadens and approaches an asymptotic value of $\Delta\kappa_g^{t,*} \approx 1.52(\Delta\kappa + \kappa)$. Note that $\Delta\kappa_g = 0$ always leads to 0 force. For small $\Delta\kappa$, $A$ becomes negative for $\Delta\kappa_g < 2(\Delta\kappa + \kappa)$. As $\Delta\kappa$ increases, $A$ becomes zero close to $\Delta\kappa_g = 2(\kappa + \Delta\kappa)$.

### Supplemental Figures

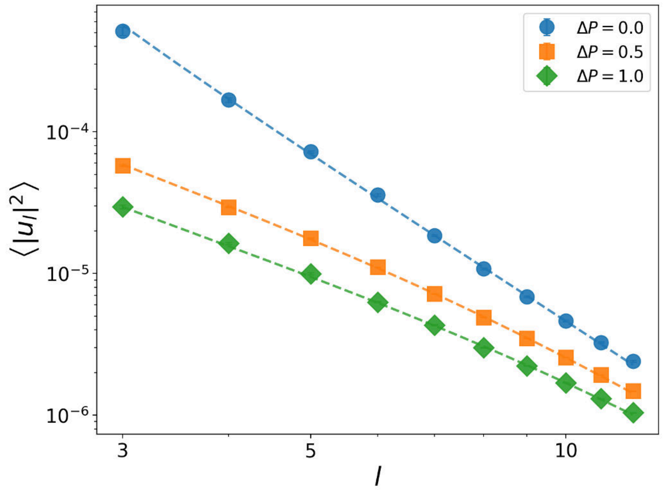


Fig. S1: Undulation spectrum of a spherical vesicle with three different values of osmotic pressure $\Delta P$. The shape of the vesicle is decomposed into spherical harmonics $Y_l^m(\theta, \phi)$, and the plot shows the mean-square amplitude $\langle |u_l|^2 \rangle$ of each mode $l$. Dashed lines are fits to the theoretical undulation spectrum of a closed spherical membrane, $\langle |u_l|^2 \rangle = \frac{k_B T}{\kappa_{\text{eff}}(l-1)(l+2)[l(l+1)+\bar{\sigma}]}$ [3], where $\kappa_{\text{eff}}$ is the effective bending rigidity, $\bar{\sigma}$ is a dimensionless membrane tension, and $l$ is the spherical harmonic mode number. Increasing osmotic pressure suppresses the amplitude of thermal fluctuations across all modes. Using Laplace's law, a pressure difference is equivalent to a surface tension $\sigma = R\Delta P/2$, which explains the suppression of fluctuations with increasing $\Delta P$.

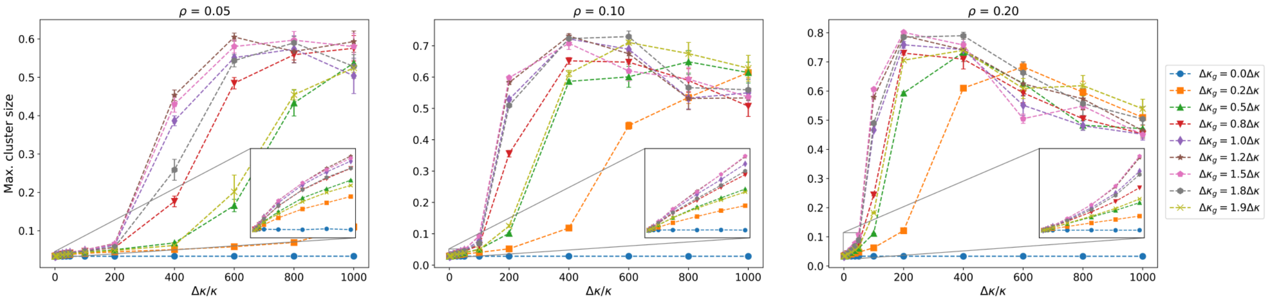


Fig. S2: Average maximum cluster size as a function of normalized inclusion bending rigidity $\Delta\kappa/\kappa$ for different normalized Gaussian moduli $\Delta\kappa_g/\Delta\kappa$. The maximum cluster size is expressed as the fraction of inclusion vertices belonging to the largest cluster in the system. Three different surface coverages, $\rho = 0.05, 0.1, 0.2$, are shown in the left, middle, and right panels, respectively.

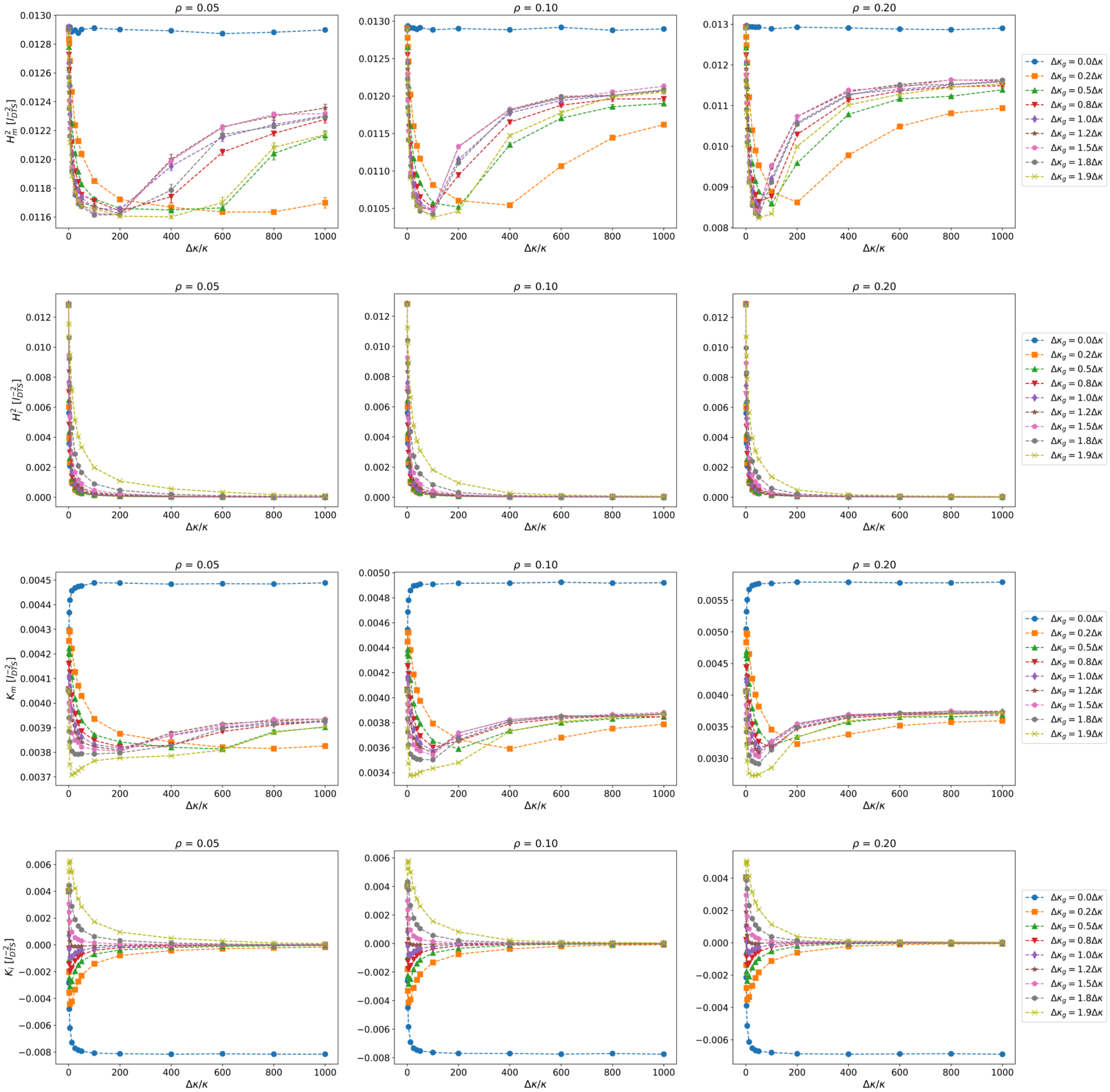


Fig. S3: Average curvature fluctuations for vertices that host an inclusion and for bare membrane vertices as a function of normalized inclusion bending rigidity $\Delta\kappa/\kappa$ for different normalized inclusion Gaussian moduli $\Delta\kappa_g/\Delta\kappa$ in a flat geometry. Each row corresponds to a different curvature measure, and each column corresponds to a different surface coverage, $\rho = 0.05, 0.1, 0.2$ (left to right). From top to bottom, the rows show: (1) average squared mean curvature for membrane particles $H_m^2$ $[l_{dts}^{-2}]$, (2) average squared mean curvature for inclusion particles $H_i^2$ $[l_{dts}^{-2}]$, (3) average Gaussian curvature for membrane particles $K_m$ $[l_{dts}^{-2}]$, and (4) average Gaussian curvature for inclusion particles $K_i$ $[l_{dts}^{-2}]$ . All quantities are normalized by the number of vertices.

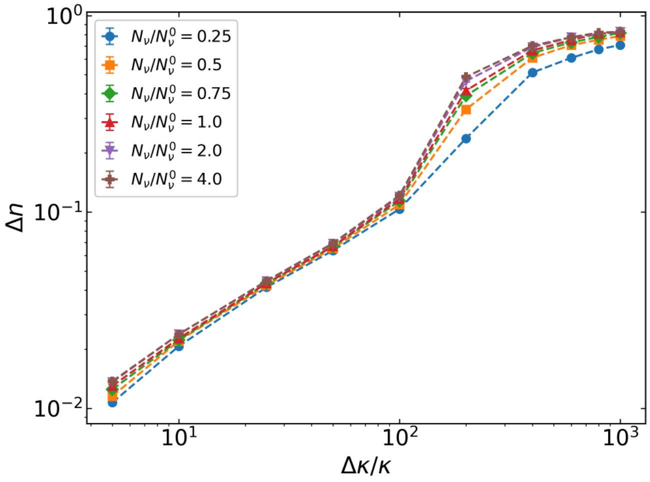


Fig. S4: Average relative inclusion-inclusion contact measure $\Delta n$ as a function of normalized inclusion bending rigidity $\Delta\kappa/\kappa$ for different system sizes $N_\nu/N_\nu^0$ ranging from 0.25–4.0. In here, $N_\nu^0$ corresponds to 1968 vertices. In the small stiffness regime, curves collapse independently of system size, consistent with a small correlation length $\xi$. Near the crossover bending rigidity, $\Delta\kappa^*$, curves systematically separate, which is the expected signature of finite-size scaling at a critical point or sharp, crossover regime, where $\xi$ is comparable to the system size $L$. At large $\xi$, curves collapse again as the system enters an ordered phase. Taken together, the system-size dependence is not a limitation of the chosen system size, but confirmatory for a large correlation length close to the crossover bending rigidity $\Delta\kappa^*$ .

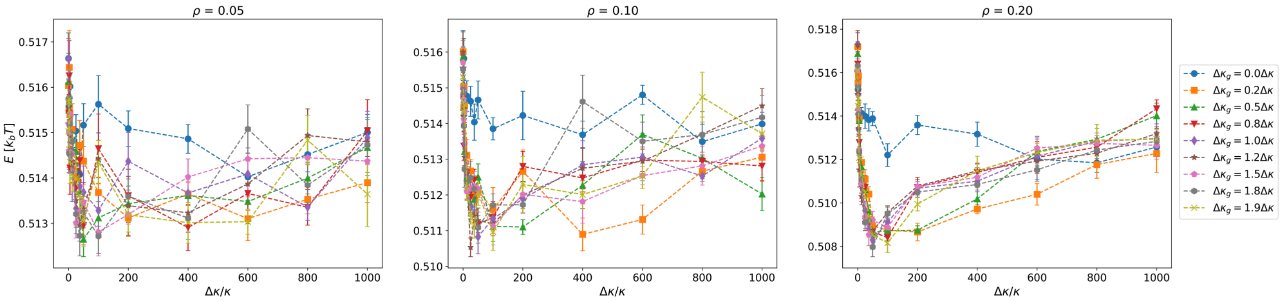


Fig. S5: Average total energy $E$ (in units of $k_bT$) per vertex as a function of normalized bending rigidity $\Delta\kappa/\kappa$ for different normalized Gaussian moduli $\Delta\kappa_g/\Delta\kappa$. Three different surface coverages, $\rho = 0.05, 0.1, 0.2$, are shown in the left, middle, and right panels, respectively.

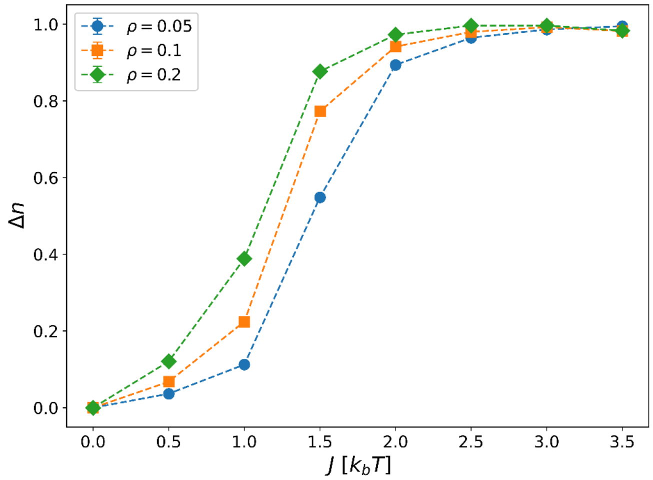


Fig. S6: Average relative inclusion-inclusion contact measure $\Delta n$ as a function of interaction constant $J$ (in units of $k_b T$) for three different inclusion surface coverages.

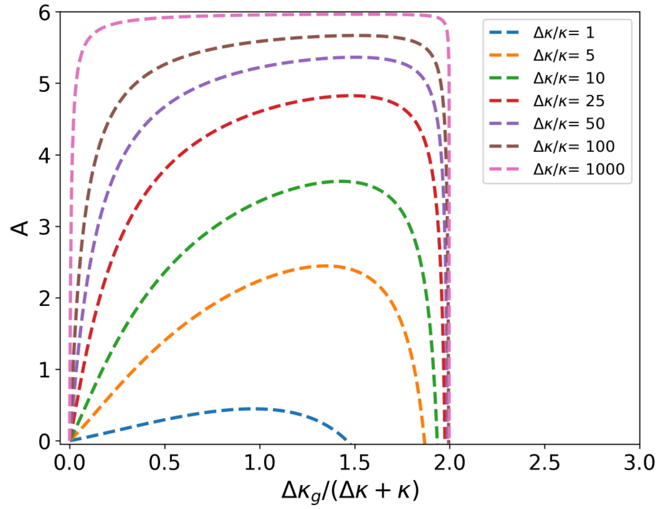


Fig. S7: Prefactor for the two-body theoretical force as a function of normalized inclusion Gaussian modulus $\Delta\kappa_g/\Delta\kappa$ for different normalized inclusion bending rigidities $\Delta\kappa/\kappa$.

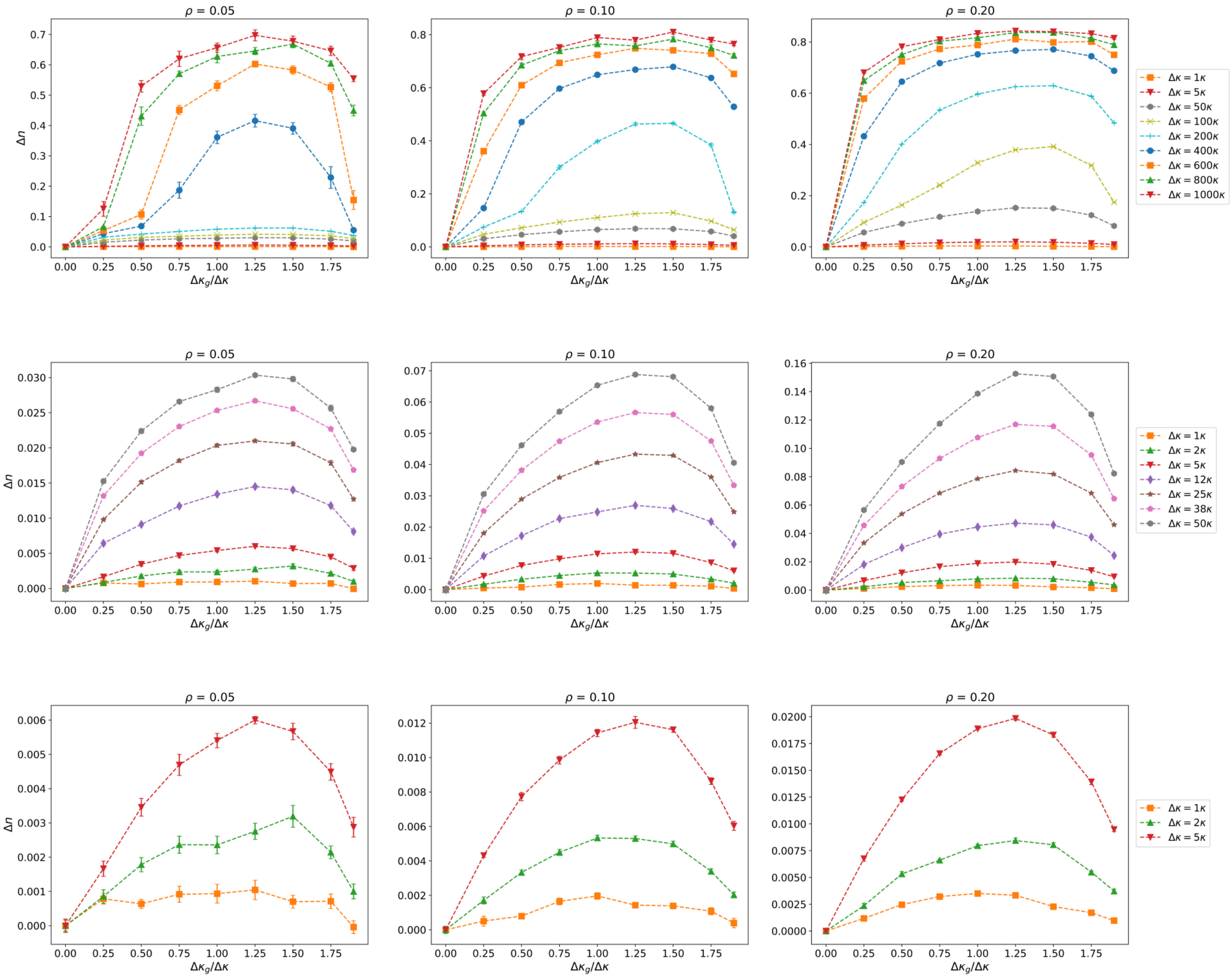


Fig. S8: Average relative inclusion-inclusion contact measure $\Delta n$ as a function of normalized Gaussian modulus $\Delta\kappa_g/\Delta\kappa$ for different inclusion bending rigidities $\Delta\kappa/\kappa$. Each column corresponds to a different surface coverage, $\rho = 0.05, 0.1, 0.2$ (left to right). In the top row, all inclusion bending rigidities are shown; however, lower bending rigidities appear nearly flat when plotted against $\Delta\kappa_g/\Delta\kappa$ due to the large range of the y-axis. For this reason, the middle and bottom rows display subsets of smaller inclusion bending rigidities for better visibility.

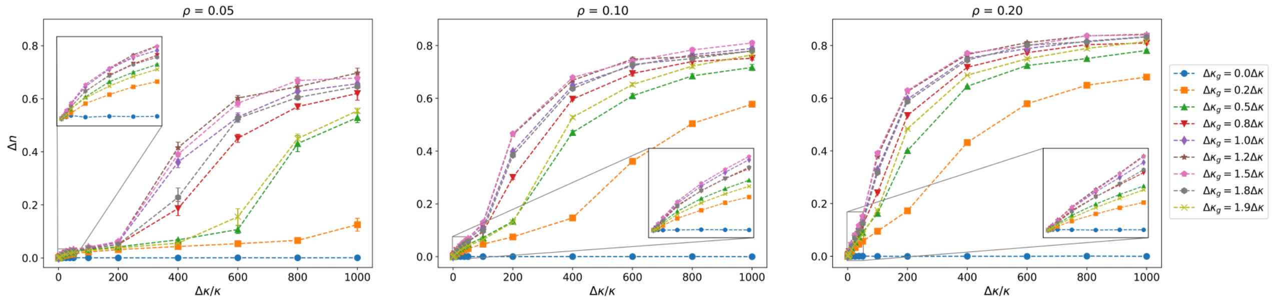


Fig. S9: Average relative inclusion-inclusion contact measure $\Delta n$, as a function of normalized inclusion bending rigidity $\Delta\kappa/\kappa$ for different normalized Gaussian moduli $\Delta\kappa_g/\Delta\kappa$. Three surface coverages $\rho = 0.05, 0.1, 0.2$, are shown in the left, middle, and right panels, respectively.

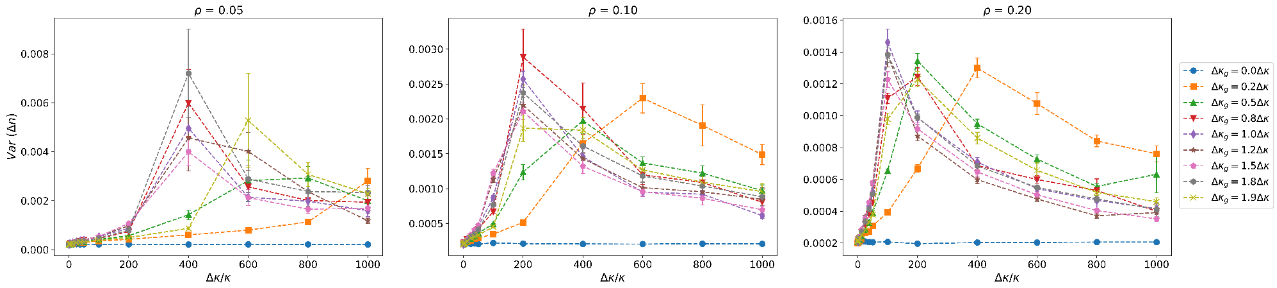


Fig. S10: Variance of the inclusion-inclusion contact measure, $Var(\Delta n) = \langle \Delta n^2 \rangle - \langle \Delta n \rangle^2$, as a function of normalized inclusion bending rigidity $\Delta\kappa/\kappa$ for different normalized Gaussian moduli $\Delta\kappa_g/\Delta\kappa$. Three surface coverages $\rho = 0.05, 0.1, 0.2$, are shown in the left, middle, and right panels, respectively.

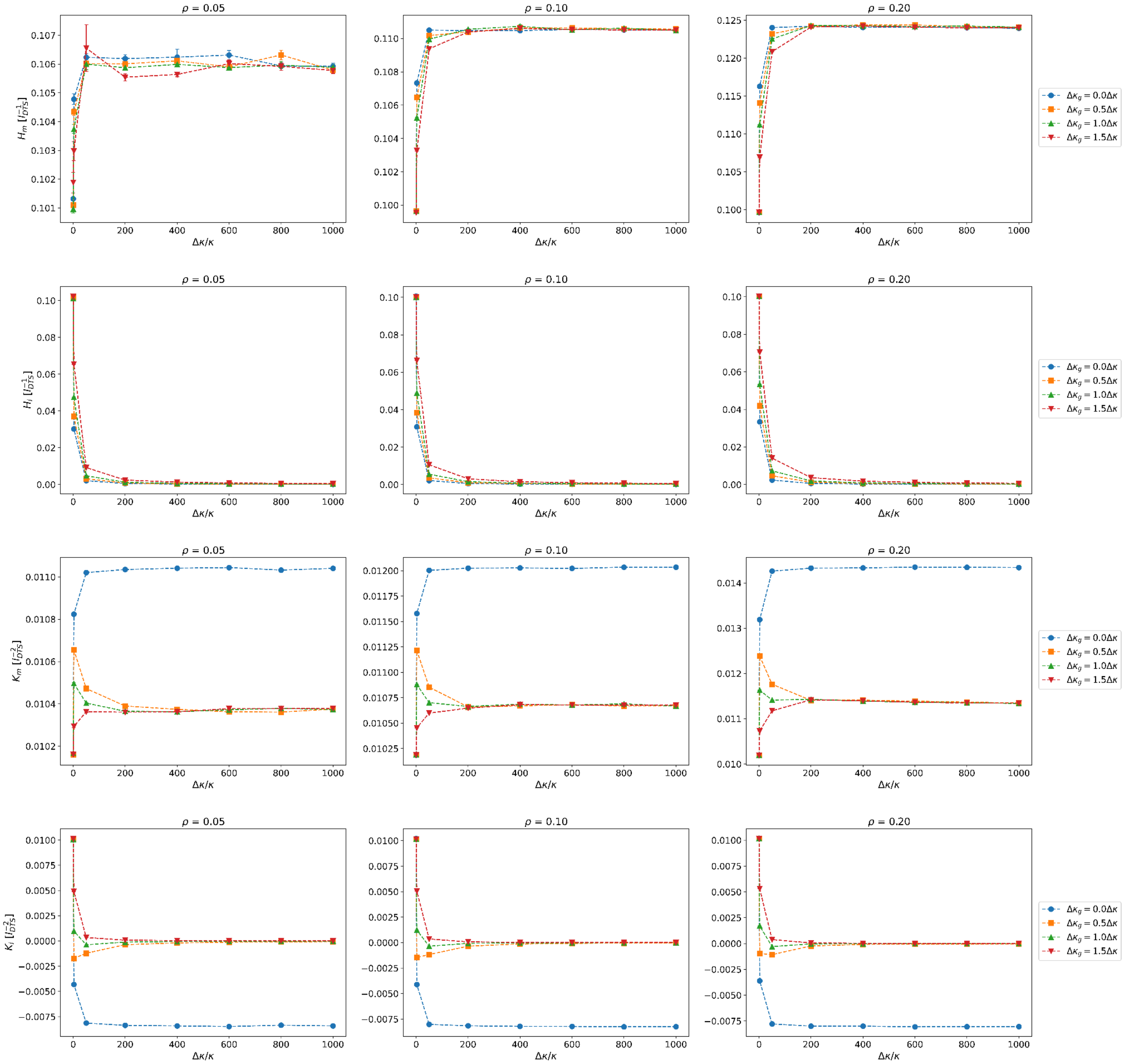


Fig. S11: Average curvature fluctuations for vertices that host an inclusion and for bare membrane vertices as a function of normalized inclusion bending rigidity $\Delta\kappa/\kappa$ for different normalized inclusion Gaussian moduli $\Delta\kappa_g/\Delta\kappa$ in a spherical geometry. Each row corresponds to a different curvature measure, and each column corresponds to a different surface coverage, $\rho = 0.05, 0.1, 0.2$ (left to right). From top to bottom, the rows show: (1) average mean curvature for membrane particles $H_m$ $[l_{dts}^{-1}]$, (2) average mean curvature for inclusion particles $H_i$ $[l_{dts}^{-1}]$, (3) average Gaussian curvature for membrane particles $K_m$ $[l_{dts}^{-2}]$, and (4) average Gaussian curvature for inclusion particles $K_i$ $[l_{dts}^{-2}]$ . All quantities are normalized by the number of vertices.

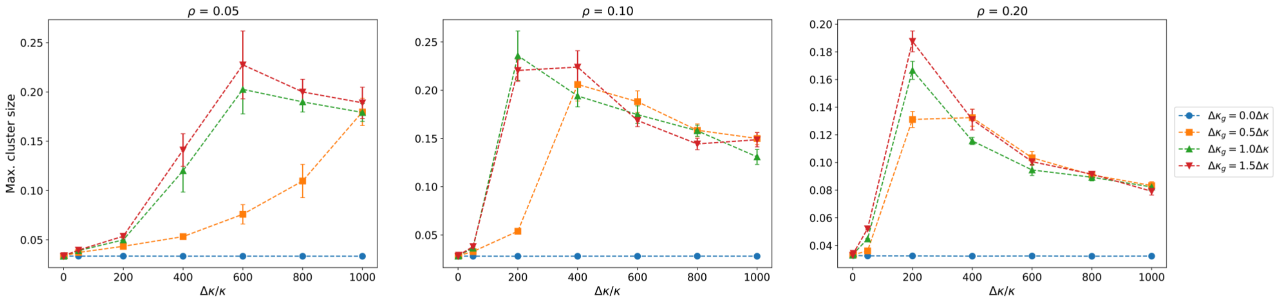


Fig. S12: Average maximum cluster size as a function of normalized inclusion bending rigidity $\Delta\kappa/\kappa$ for different normalized Gaussian moduli $\Delta\kappa_g/\Delta\kappa$ in spherical geometry. The maximum cluster size is measured as the fraction of inclusion vertices belonging to the largest cluster. Three surface coverages, $\rho = 0.05, 0.1, 0.2$, are shown from left to right.

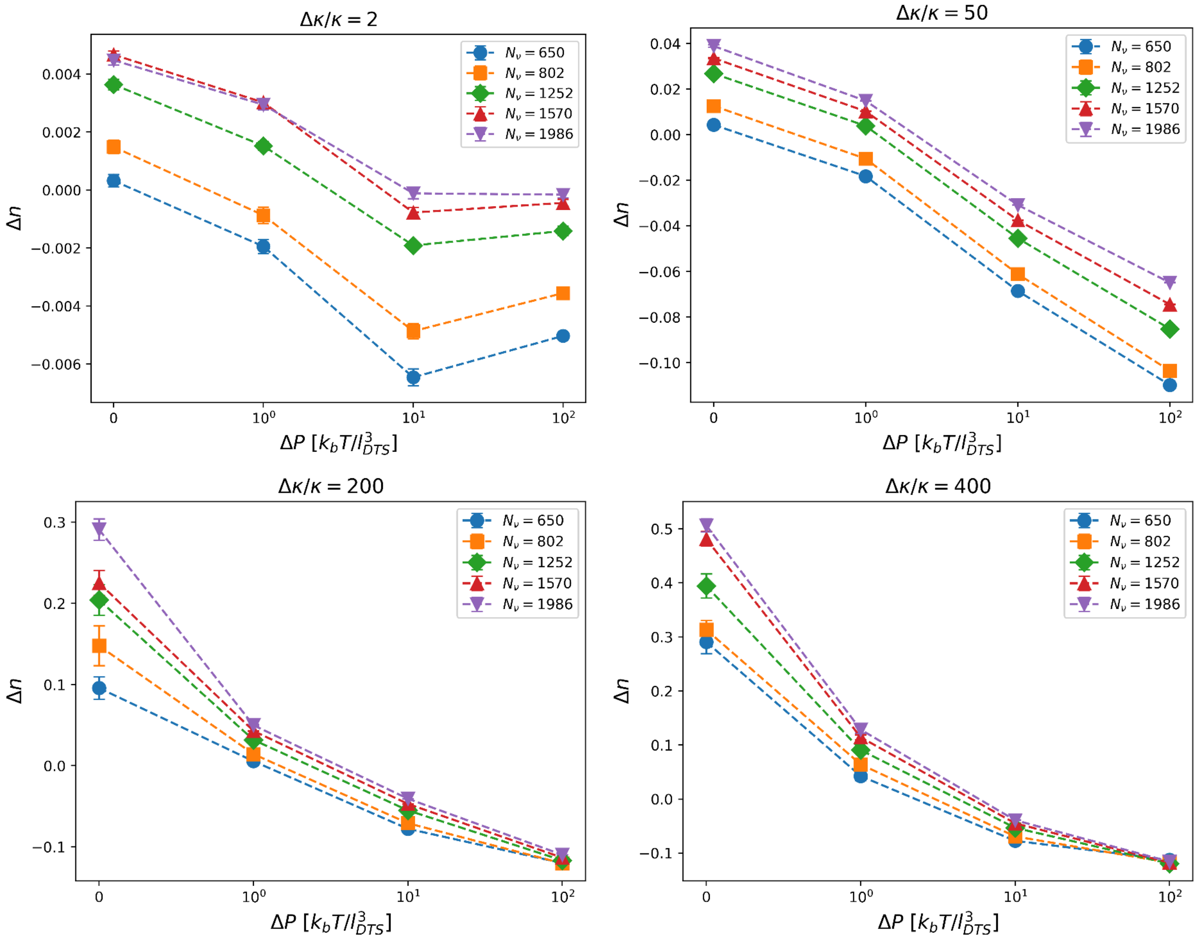


Fig. S13: Average relative inclusion-inclusion contact measure $\Delta n$ as a function of osmotic pressure $\Delta P$, in units of $k_BT/l^3_{\mathrm{DTS}}$ for vesicles of different sizes $N_\nu$. The figure is arranged as a $2 \times 2$ panel, with each subplot corresponding to a different normalized inclusion bending rigidity $\Delta\kappa/\kappa$ (indicated at the top of each panel). The inclusion Gaussian modulus is set to $\Delta\kappa_g = \Delta\kappa$, and the surface coverage is fixed at $\rho = 0.1$.

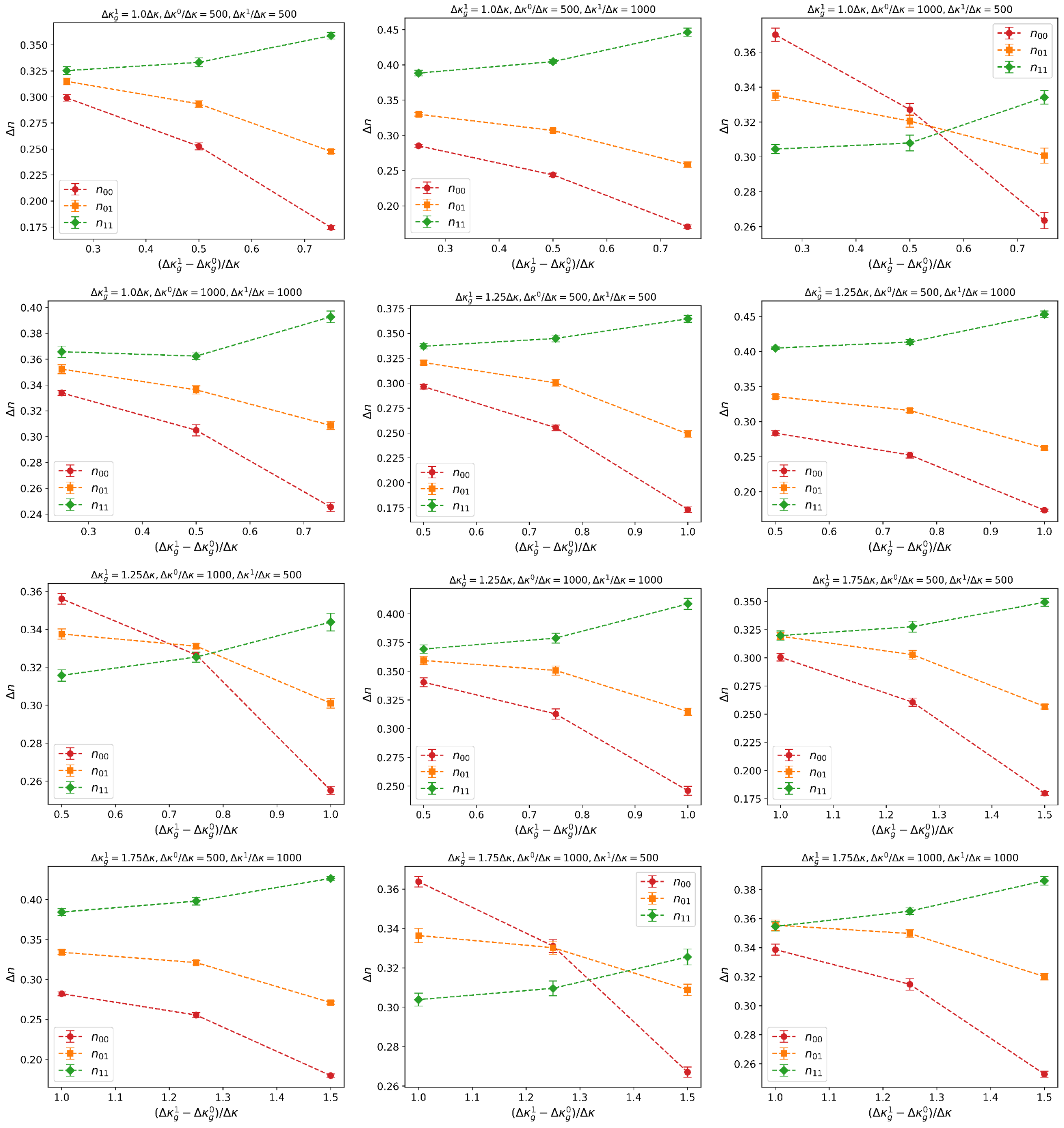


Fig. S14: Average relative inclusion-inclusion contact measure $\Delta n$ as a function of the difference in Gaussian modulus $(\Delta\kappa_g^1 - \Delta\kappa_g^0)/\Delta\kappa$ between two inclusion types. The quantities $n_{00}$ ($n_{11}$) correspond to the $\Delta n$ for inclusions of type 0 (1), while $n_{01}$ measures the $\Delta n$ between inclusions of type 0 and type 1. Each subplot is labeled with the Gaussian modulus of type 1 inclusions, $\Delta\kappa_g^1$, together with the normalized inclusion bending rigidities of inclusions of type 0 and 1, $\Delta\kappa^i/\kappa$ ($i = 0, 1$). The surface coverage of both inclusion types is fixed at $\rho_0 = \rho_1 = 0.05$. The figure is arranged as a $4 \times 3$ panel layout.

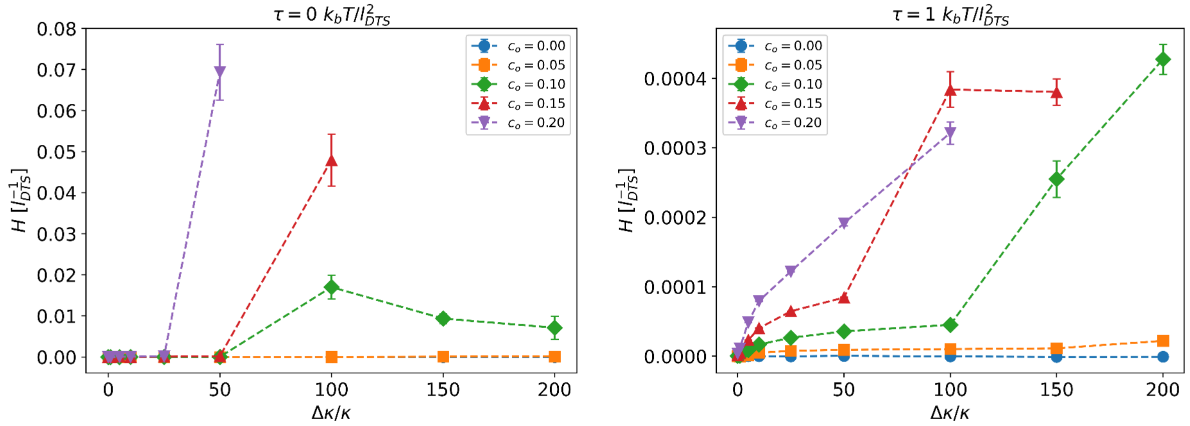


Fig. S15: Average normalized total mean curvature $H$ (in units of $l_{\rm DTS}^{-1}$) as a function of inclusion bending rigidity $\Delta\kappa/\kappa$ for different inclusion local curvatures $c_0$ (in units of $l_{\rm DTS}^{-1}$). The inclusion Gaussian modulus is fixed at $\Delta\kappa_g = \Delta\kappa$, and the surface coverage is set to $\rho = 0.1$.

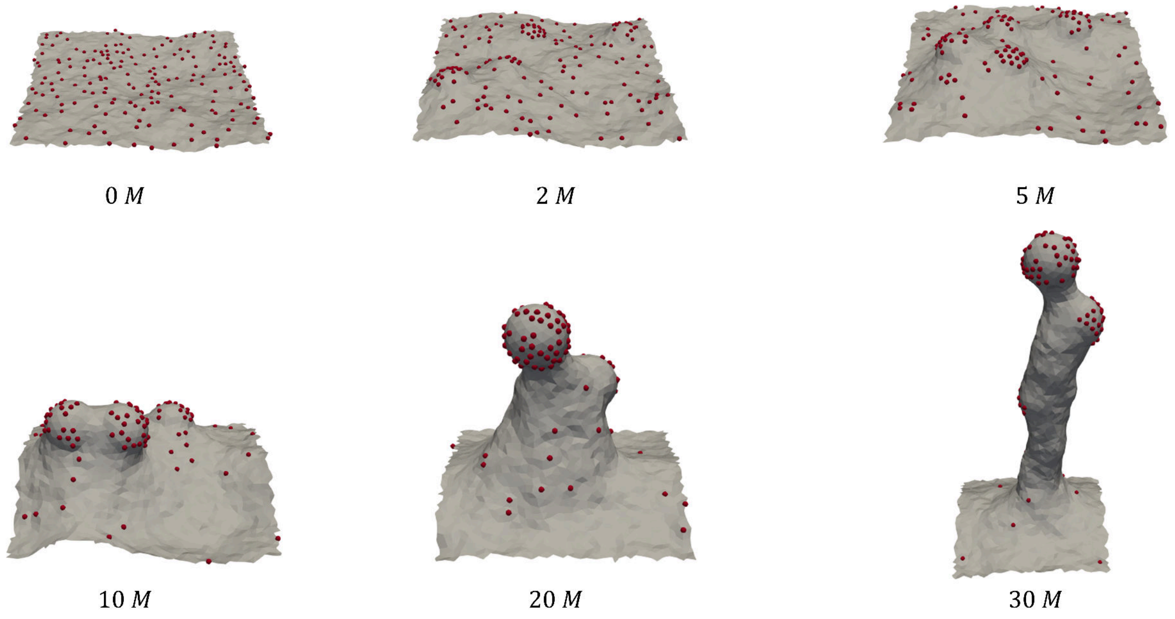


Fig. S16: Simulations snapshots corresponding to a membrane partially covered by curvature-inducing, stiff inclusions. Below the frame, the number of MC moves is shown in million MC steps units (M). The membrane bending rigidity is set to $\kappa = 10\ k_bT$. Inclusions interact with the membrane with $\Delta\kappa = 50\ \kappa$, $\Delta\kappa_g = \Delta\kappa$ and $c_o = 0.2\ l_{dts}^{-1}$. The inclusion surface coverage is $\rho = 0.1$.

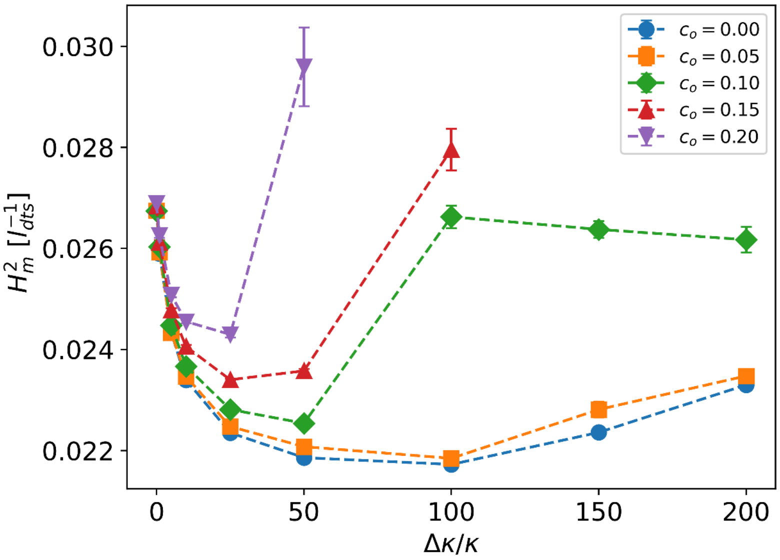


Fig. S17: Average squared mean curvature for membrane particles $H_m^2$ (in units of $l_{dts}^{-2}$) as a function of inclusion bending rigidity $\Delta\kappa/\kappa$ for different inclusion local curvatures $c_0$ (in units of $l_{\mathrm{DTS}}^{-1}$). The inclusion Gaussian modulus is fixed at $\Delta\kappa_g = \Delta\kappa$, and the surface coverage is set to $\rho = 0.1$. Surface tension is set to $\tau = 0\ k_bT/l_{dts}^2$

.
